# Dynamical BLUP modeling of reaction norm evolution, accommodating changing environments, overlapping generations, and multivariate data

**DOI:** 10.1101/2023.04.09.536146

**Authors:** Rolf Ergon

**Affiliations:** University of South-Eastern Norway, Porsgrunn, Norway

**Keywords:** Dynamical BLUP, Mean reaction norm parameter updating, Microevolution vs. plasticity disentanglement, Overlapping generations, Robertson’s secondary theorem of natural selection, Random effect variance estimation

## Abstract

For theoretical studies, reaction norm evolution in a changing environment can be modeled by means of the multivariate breeder’s equation, with the reaction norm parameters treated as traits in their own right. This is, however, not a feasible approach for use of field data, where the intercept and slope values are not available. An alternative approach is to use infinite-dimensional characters and smooth covariance function estimates found by, i.e., random regression. This is difficult because of the need to find for example polynomial basis functions that fit the data reasonably well over time, and because reaction norms in multivariate cases are correlated, such that they cannot be modeled independently.

Here, I present an alternative approach based on a multivariate linear mixed model of any order, with dynamical incidence and residual covariance matrices that reflect the changing environment. From such a mixed model follows a dynamical BLUP model for prediction of the individual reaction norm parameter values at any given parent generation, and for updating of the mean reaction norm parameter values from generation to generation by means of Robertson’s secondary theorem of natural selection. This will, e.g., make it possible to disentangle the microevolutionary and plasticity components in climate change responses. The BLUP model incorporates the additive genetic relationship matrix in the usual way, and overlapping generations can easily be accommodated.

Parameters are assumed to be known and constant, but it is discussed how they can be estimated by means of a prediction error method. The identifiability by use of field or laboratory data containing environmental, phenotypic, fitness, and additive genetic relationship data, is an important feature of the proposed model.

## 1. Introduction

A reaction norm describes the phenotypes that a genotype can produce across a range of environments. Mean reaction norms in a population can evolve, and this evolution can be modeled by use of several methods. In its simplest form a mean reaction norm is characterized by an intercept value and a plasticity slope value, but it may also be natural to use models with a multiple of reaction norms, and they may be non-linear.

A common model is the multivariate breeder’s equation (Lande, 1979), where the mean reaction norm parameters may be treated as traits in their own right, as in, e.g., Lande (2009). For applications on field data from studies of wild populations, there are two problems with such models. First, the individual reaction norm parameters are not available, and second, additive genetic relationships in the population cannot be taken into account. The first problem can be solved by a linear transformation as shown in Ergon (2022a), and as used for comparison purposes in Section 2, but the second problem will still exist.

An alternative approach for reaction norm modeling is to use infinite-dimensional characters, as in a method introduced by Kirkpatrick and Heckman (1989), and as applied on reaction norms by Gomulkiewicz and Kirkpatrick (1992). It is then necessary to obtain smooth covariance function estimates (Kingsolver et al, 2001), and one method for that purpose is random regression (Shaeffer, 2004), where individual breeding values are modeled as relatively simple weighted sums of basis functions. Additive genetic relationship matrices can be included in such models (Oliviera et al., 2019). An obvious difficulty is here the need to find for example polynomial basis functions that fit the data reasonably well over time.

Another difficulty is that the reaction norms in multivariate cases are correlated, such that they cannot be modeled independently.

In the present article I introduce a multivariate modeling approach based on best linear unbiased predictions (BLUP), allowing for multiple (and potentially correlated) traits to have joint norms of reaction, which can be linear or approximated by power series (Gavrilets and Scheiner, 1993). This is a dynamical BLUP model in the sense that the incidence matrix for the random effects and the residual covariance matrix are functions of the changing environment. I will develop the theory under the assumption that the parameters in the model are known, but as will be shown separately they can also be identified by a prediction error method, as introduced in a microevolutionary context in Ergon (2022a,b). This identification aspect is an essential prerequisite, i.e., the model must be identifiable from available environmental, phenotypic, fitness, and additive genetic relationship data.

Best linear unbiased predictions (BLUP) utilizing linear mixed models with fixed and random effects, are extensively used in domestic animal and plant breeding (Robinson, 1991; Ch. 26, Lynch and Walsh, 1998; Arnold et al., 2019). These methods may also be applied on wild populations (Kruuk, 2004; Nussey et al., 2007), although such uses have been criticized owing to errors in estimated variances of the random effects (Hadfield et al., 2010). An important application is the disentanglement of microevolutionary and plasticity components in for example climate change responses (Merilä and Hendry, 2014; Ergon, 2022a,b). The basic BLUP equations was first developed in summation form (Henderson, 1950), but as done here it is more convenient to use matrix formulations.

This article will be focused on how mean reaction norm parameter values, and thus mean phenotypic traits, evolve under the influence of environmental cues and changes of the fitness landscape. Such evolution of reaction norms and phenotypic traits seeks to maximize the mean fitness of a given population, and changes in the location of fitness peaks in the phenotypic space are therefore the driving force. I will thus study the dynamics of microevolutionary systems, mainly by use of BLUP, but also with reference to the well-known multivariate breeder’s equation (Lande, 1979). Although fitness can be defined as the long-run growth rate (Sæther and Engan, 2015), or for non-overlapping generations the expected geometric mean fitness (Autzen and Okasha, 2022), I will in simulations simply use the number of surviving descendants as a measure of individual fitness (Ch. 6, Rice, 2004).

It is well known that BLUP underestimates the variances of the random effects in linear mixed models (Ch. 26, Lynch and Walsh, 1998; Hadfield et al., 2010). Here, I will show why and to which extent that is necessary in order to obtain the correct incremental changes in mean reaction norm parameter values from generation to generation. I will also show that these changes may be found from Robertson’s secondary theorem of natural selection (Robertson, 1966) applied on the predicted random effects. This also applies to non-plastic organisms, where the mean reaction norms degenerate into mean phenotypic trait values (Ergon, 2022c).

The dynamical BLUP model with Robertson updating of mean reaction norm parameter values, makes use of the additive genetic relationship matrix ***A***_*t*_ in a standard way (Ch. 26, Lynch and Walsh 1998). The theoretical treatment is limited to cases where only mean phenotypic traits are included in the fixed effects, and for simplicity it assumes that generations are non-overlapping. It is, however, also shown how cases with overlapping generations can be handled in a straightforward way. For cases with sexual reproduction, I assume a hypothetical single parent (mid-parent) occupying an intermediate phenotypic position between the two parents (Ch. 7, Rice, 2004).

The theoretical development will be general, i.e., for any number of phenotypic traits and any number of environmental cues in the model. For clarity of presentation, however, some details will be given for a system with only two phenotypic traits and two environmental cues. A similar limited system will also be used in simulations.

The additive genetic and phenotypic covariance matrices, ***G*** and ***P***, are here assumed to be constant and known. They may, however, be estimated by means of a prediction error method (PEM), utilizing information contained in environmental cues and individual phenotypic trait values over many generations, as well as fitness information (Ergon, 2022a,b). With plastic traits in the dynamical BLUP model, restricted maximum likelihood (REML) methods applied on data from a single generation cannot be used for this purpose. The simple reason for this is that each element in the residual covariance matrix is a function of several non-additive effects, such that the REML equations become indeterminate.

As developed theoretically, and verified in simulations, the dynamical BLUP model with an additive genetic relationship matrix equal to an identity matrix, i.e., with random mating in an unbred population, will give the same results as a selection gradient prediction method (GRAD) based on the multivariate breeder’s equation (Ergon, 2022a,b). For large populations, these results will asymptotically also be the same as from the multivariate breeder’s equation directly.

After this introduction, Theory and Methods follow in Section 2, Simulations in Section 3, and Summary and Discussion in Section 4. Proofs of two theorems are given in Appendices A and B. For the interested reader, a user guide is given in Appendix C, including the procedure for PEM system identification. MATLAB code for the simulations is given in Supplementary Material.

## 2. Theory and Methods

### 2.1 Notation

Mathematical symbols with descriptions in the order they appear in equations are shown in Table 1.

**Table 1.** Mathematical symbols with description

| Symbol | Description |
| --- | --- |
| $\Delta\bar{z}_t$ | Incremental change in mean trait from generation $t$ to generation $t + 1$ |
| $w_{i,t}$ | Relative individual fitness, $w_{i,t} = W_{i,t}/\bar{W}_t$ |
| $z_{i,t}, y_{i,t}, \mathbf{z}_{i,t}, \mathbf{y}_{i,t}$ | Individual traits, and vectors of individual traits |
| $\mathbf{G}$ | Additive genetic covariance matrix, with block elements $\mathbf{G}_{aa}, \mathbf{G}_{ab}, \mathbf{G}_{bb}$ |
| $\mathbf{P}$ | Phenotypic covariance matrix, with block elements $\mathbf{P}_{aa}, \mathbf{P}_{ab}, \mathbf{P}_{bb}$ |
| $\boldsymbol{\beta}_t$ | Selection gradient |
| $a'_{i,t} + v_{i,t}$ | Individual intercept deviations around mean values $\bar{a}_t$ and 0, respectively, with additive and non-additive effects |
| $b'_{i,t} + \eta_{i,t}$ | Individual plasticity slope deviations around mean values $\bar{b}_t$ and 0, respectively, with additive and non-additive terms |
| $\Delta\bar{a}_t, \Delta\bar{b}_t$ | Incremental changes in mean reaction norm parameter values |
| $\sigma_v^2, \sigma_\eta^2$ | Variances of non-additive effects |
| $u_t, \mathbf{u}_t$ | Environmental variable, and vector of environmental variables |
| $P_{yy}$ | Variance of $y_{i,t}$ |
| $\mathbf{X}$ | Design matrix in linear mixed model |
| $\tilde{\mathbf{Z}}_t = \mathbf{Z}_t \otimes \mathbf{I}_n$ | Incidence matrix in linear mixed model, with $\mathbf{Z}_t = f(\mathbf{u}_t)$ |
| $\mathbf{x}_t$ | Random effects in linear mixed model, for special case with 2 traits and 2 environm. variables, $\mathbf{x}_t = [\mathbf{a}'_{1,t} \quad \mathbf{a}'_{2,t} \quad \mathbf{b}'_{11,t} \quad \mathbf{b}'_{12,t} \quad \mathbf{b}'_{21,t} \quad \mathbf{b}'_{22,t}]^T$ |
| $\mathbf{e}_t$ | Residual vector in mixed model, with $\mathbf{e}_t = f(\mathbf{u}_t)$ |
| $\mathbf{A}_t$ | Additive genetic relationship matrix |
| $\tilde{\mathbf{G}}_t = \mathbf{G} \otimes \mathbf{A}_t$ | Kronecker covariance matrix for BLUP model |
| $\tilde{\mathbf{R}}_t = \mathbf{R}_t \otimes \mathbf{I}_n$ | Kronecker residual matrix for BLUP model, with $\mathbf{R}_t = E[\mathbf{e}_t \mathbf{e}_t^T]$ |
| $\hat{z}'_{i,t}$ | Predicted individual intercept or slope deviations around mean values |
| $\bar{z}_t^{parents}, \bar{z}_t^{offspring}$ | Mean reaction norm parameter values for parents and offspring |

### 2.2 Introductory example

For a simple toy example, intended to ease the readers into the concepts used below, consider a single trait *y*_*i,t*_ measured on a single individual. In this case, the BLUP estimate of the true additive genetic value for that individual is 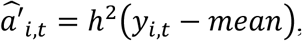, where *h*^2^ is the heritability, while *mean* denotes any fixed-effects adjustment. If we substitute this estimate into Robertson’s secondary theorem of natural selection (Ch. 6, Walsh and Lynch, 2018), we find the between-generation response 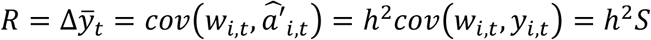, where *w*_*i,t*_ is the relative fitness, while *S* is the Robertson-Price within-generation change in the mean. In this way we recover the standard univariate breeder’s equation.

What is done below is to consider a much more complicated phenotype (an observed vector *y*_*t*_ of individual focal traits with reaction norms) and use BLUP to estimate the vector of additive effects associated with the norm of reaction functions, with these BLUPs then substituted into the expression for Robertson’s secondary theorem of natural selection. The incremental changes in the mean reaction norm parameter values thus follow from 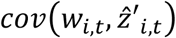, where 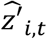, stands for the BLUP estimates of the the true additive genetic values involved (Eq. (11) below).

For the special case with an additive genetic relationship matrix ***A***_*t*_ = ***I***_*n*_, the BLUP estimates yield a matrix-based inheritance expression (using the correlated nature of the random effects) to replace *h*^2^, and a Robertson-Price term *cov*(*w*_*i,t*_, *y*_*i,t*_) to measure phenotypic selection (Eq. (12) below). From a practical point of view, Eq. (12) is unnecessary, but it is included for the purpose of comparisons with results from the multivariate breeder’s equation.

### 2.3 Background theory

For the development of the dynamical BLUP matrix equation that follows, we need some background theory. First, the Price equation for selection in a population with *n* individuals says that the evolution of the mean trait of an *n* × 1 vector ***z***_*t*_ of individual quantitative traits is described by (Price, 1970, 1972)

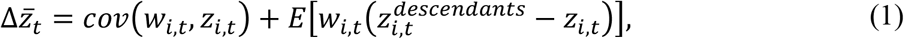

where 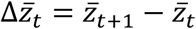 is the incremental change in mean trait value from generation to generation, and where *z*_*i,t*_ is an individual trait. Here, *w*_*i,t*_ is the relative individual fitness, i.e., individual fitness divided by the mean fitness in the population, while 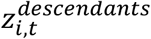 is the mean trait of the descendants (and the parent if it survives) of individual *i* in generation *t*. The trait *z*_*i,t*_ may be any property we can assign a numerical value to, not necessarily biological. In a biological context the trait may be a behavioral, morphological, or physiological characteristic, but it may also be a parameter in a reaction norm model that describes a plastic organism. Disregarding the second term on the righthand side of Eq. (1), we find the Robertson-Price identity 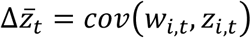 (Robertson, 1966; Ch. 6, Walsh and Lynch, 2018), as referred to above, and which we will use below.

Second, we need to see how the multivariate breeder’s equation (Lande, 1979; Lande and Arnold, 1983),

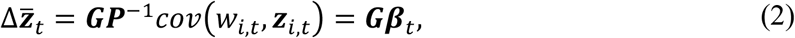

where *β*_*t*_ is the selection gradient, can be applied on the parameters in a reaction norm model. Eq. (2) was derived from a multivariate version of Eq. (1), which requires several assumptions, as detailed in Ergon (2019,2022c):

1. The vector ***z***_*i,t*_ of individual phenotypic traits is the sum of independent additive genetic effects ***x***_*i,t*_ and non-additive environmental and genetic effects ***e***_*i,t*_, i.e., ***z***_*i,t*_ = ***x***_*i,t*_ + ***e***_*i,t*_.
2. The non-additive effects ***e***_*i,t*_ are zero mean, independent and identically distributed (iid) random variables.
3. There are no expected fitness weighted changes in the individual additive genetic effects ***x***_*i,t*_ from one generation to the next besides selection, i.e., 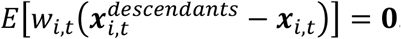.
4. The additive genetic effects ***x***_*i,t*_, and the environmental effects ***e***_*i,t*_ and 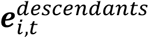, are multivariate normal.
5. The additive genetic effects ***x***_*i,t*_ and non-additive effects ***e***_*i,t*_ and 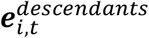 influence individual fitness only through ***z***_*i,t*_.
6. All individuals in the population are genetically unrelated, which means that the additive genetic relationship matrix ***A***_*t*_ is a unity matrix.

In what follows, we will make use of Assumptions 1, 2, and 3, while Assumptions 4, 5, and 6 will be used only indirectly when the BLUP results with ***A***_*t*_ = ***I***_*n*_ are compared with results based on the multivariate breeder’s equation.

In order to see how Eq. (2) can be applied on the parameters in a reaction norm model, we may use an individual intercept-slope model based on Assumptions 1 and 2 above (Lande, 2009),

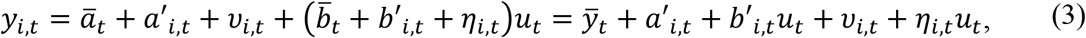

where *u*_*t*_ is an environmental cue, and where the mean reaction norm is _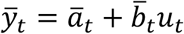_. Here, 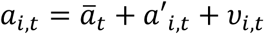 and 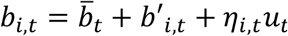 are the individual intercept and slope parameters, where *ν*_*i,t*_ and *η*_*i,t*_ are iid and zero mean random variables, with variances 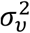 and 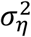, respectively. We thus use 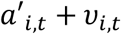 and 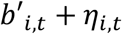 to denote individual deviations from mean values 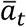 and 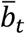, respectively, where 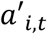 and 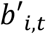 are the additive genetic components of these deviations. Such additive genetic deviations will be the random effects in the linear mixed model developed below, and thus the random effects that are predicted by use of the BLUP equations.

When the reaction norm parameters in Eq. (3) are treated as quantitative traits in their own right, Eq. (2) leads to

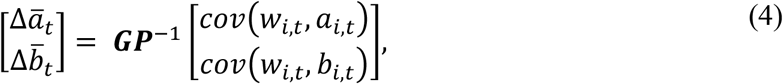

with ***G*** and ***P*** are given by 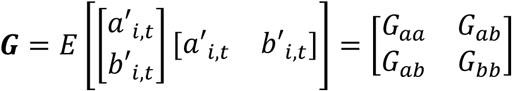 and 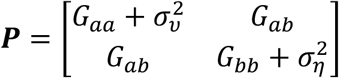.

The additive genetic and phenotypic covariance matrices ***G*** and ***P*** may by time-varying, but for simplicity we will here assume that they are constant. As discussed in Ergon (2022a), it is essential that the environmental input in Eq. (3) has a proper reference value, and for simplicity we here assume an environmental scale such that the reference environment is zero. Note that the model in Eq. (4) cannot be identified by use of available environmental, phenotypic, fitness, and additive genetic relationship data, where *a*_*i,t*_ and *b*_*i,t*_ are not included.

For comparisons with BLUP results, we finally need an identifiable version of the multivariate breeder’s equation. As shown in Ergon (2022a), Eq. (4) can by use of a linear transformation of the vector [*a*_*i,t*_ *b*_*i,t*_]^*T*^ onto the vector [*a*_*i,t*_ *b*_*i,t*_ *y*_*i,t*_]^*T*^ be reformulated into the selection gradient (GRAD) form,

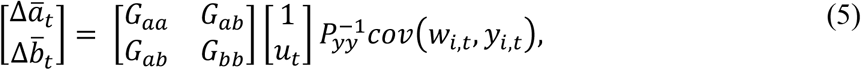

where selection with respect to *a*_*i,t*_ and *b*_*i,t*_, as in Eq. (4), is replaced by selection with respect to *y*_*i,t*_. Here, 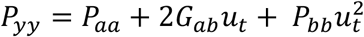, while 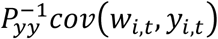 is the selection gradient.

It is essential to note that Eqs. (4) and (5) give identical results only asymptotically, when the population size *n* → ∞, and the reason for that is the differences in how the covariance functions are used in the two equations. As we will see, extensions of Eq. (5) are possible for more complex reaction norm models. This is interesting because with an additive genetic relationship matrix ***A***_*t*_ = ***I***_*n*_, the dynamical BLUP model we will develop results in incremental changes in mean reaction norm parameter values, that are identical to those found from an extended version of Eq. (5). Here, we should finally note that since Eq. (5) is derived from the multivariate breeder’s equation (2), it is valid only under Assumptions 1 to 6 above.

### 2.4 Development of the dynamical BLUP model

For clarity of presentation, some details will here be limited to a system with *p* = 2 phenotypic traits, and *q* = 2 environmental cues, and a similar simplified system will also be used in the simulations. The theory will, however, be developed in such a way that extensions to higher dimensions are obvious.

For a specific trait *j*, the individual reaction norm model with *q* = 2 environmental cues is

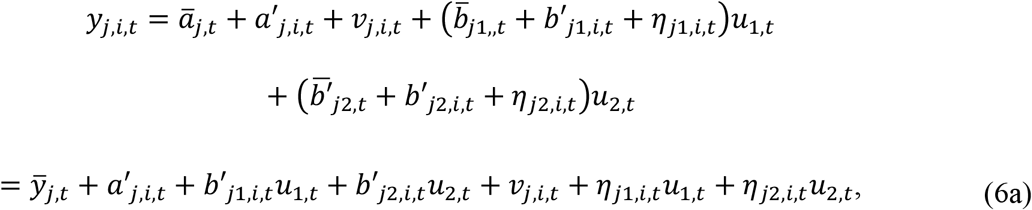

where 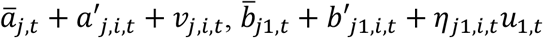 and 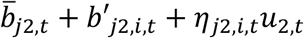 are the individual parameter values, while

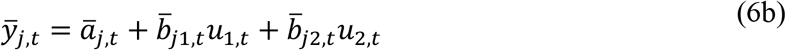

is the mean trait value. For a population with *n* individuals we may collect 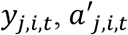, etc., in *n* × 1 vectors, and obtain the individual trait vector for trait *j*,

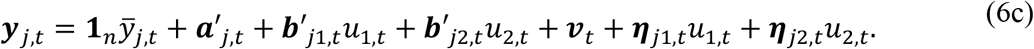

With *p* = 2 traits and *q* = 2 environmental cues, we thus obtain the linear mixed model in general form (Ch. 26, Lynch and Walsh 1998) (with 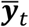 as fixed effects vector)

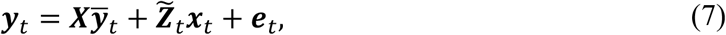

with 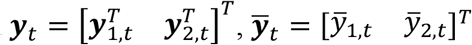, 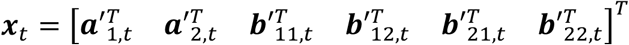, and 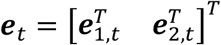, where *e*_1,*t*_ = *v*_1,*t*_ + *η*_11,,*t*_*u*_1,*t*_ + *η*_12,*t*_ *u*_2,*t*_, and *e*_2,*t*_ = *v*_2,*t*_ + *η*_21,,*t*_ *u*_1,*t*_ + *η*_22,*t*_ *u*_2,*t*_. Here, 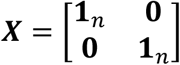, with dimension *pn* × *p* = 2*n* × 2, while 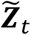 has dimension *pn* × *p*(1 + *q*)*n* = 2*n* × 6*n*. In more detail we have 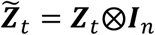, where 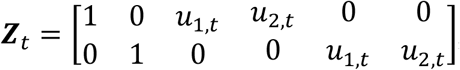, and where ⨂ is the Kronecker product operator, which means that all elements in **Z**_*t*_ should be multiplied by ***I***_*n*_. For use in Appendix B we may note that 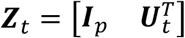, with 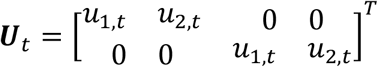. It is an essential feature of the model in Eq. (7) that 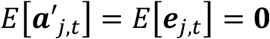, such that 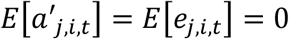 for *j* = 1 to *p*, and that 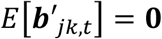, such that 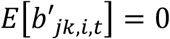 for *j* = 1 to *p* and *k* = 1 to *q* (Ch. 26, Lynch and Walsh 1998). We thus also have that 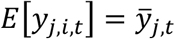.

The random effects in Eq. (7) may all be correlated, with an additive genetic covariance matrix 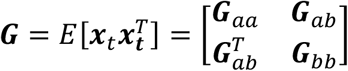. The residuals ***e***_1,*t*_ and ***e*** _2,*t*_ in Eq. (7) are assumed to be uncorrelated, with a covariance matrix 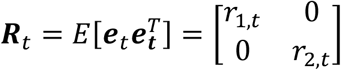, where 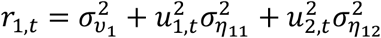 and 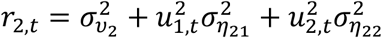 with 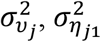 and 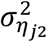 as the variances of the non-additive effects υ_*j,i,t*_, *η*_*j*1,*i,t*_ and *η*_*j*2,*i,t*_ according to Eq. (6a). Also ***G*** may in general be a function of time, but we will here assume that it is constant. We will assume that ***G*** and ***R***_*t*_ are known, although they may in practice be estimated by use of restricted maximum likelihood (REML) (Ch. 27, Lynch and Walsh 1998), or a prediction error method (Ergon, 2022a,b). Note, however, that REML equations for a single generation will be indeterminate, i.e., we can find *r*_1,*t*_ and *r*_2,*t*_, but not all the residual covariances in the expressions for *r*_1,*t*_ and *r*_2,*t*_. For use in a general multivariate BLUP equation, we also need the *n* × *n* additive genetic relationship matrix ***A***_*t*_ (Ch. 26, Lynch and Walsh, 1998), and the Kronecker covariance matrices 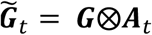 and 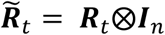.

For comparisons with predictions based on selection gradients (GRAD), as in Eq. (5), we also need the phenotypic covariance matrix 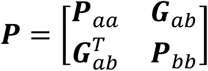, where 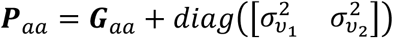 and 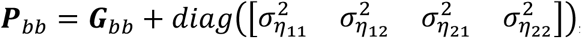, with *diag*([∙]) denoting diagonal matrices. Also ***P*** is assumed to be constant.

As will be shown in detail below, the fixed effects 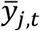 and random effects 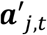 and 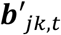 in Eq. (7) may be estimated by use of BLUP. In line with established language in the breeding literature, 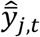 will here be called estimates, while 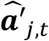 and 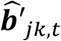 will below be called predictions. See Robinson (1991) for a discussion of the historical background for this terminology. From the mixed model (7) follows the multivariate BLUP equation in matrix form (Henderson, 1950; Ch. 26, Lynch and Walsh, 1998),

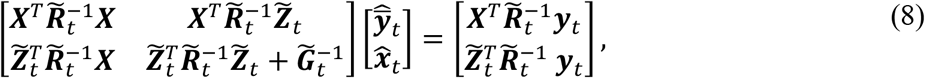

where, for *p* = 2 and 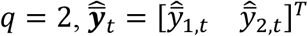 and 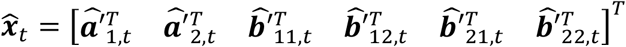. Here, 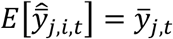 and 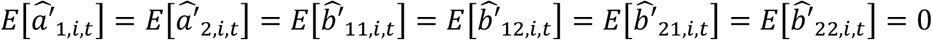. Note that the derivation of Eq. (8) does not necessarily require an assumption of normal data (Robinson 1991).

### 2.5 Updating of mean reaction norm parameter values

When Eq. (8) is applied on any given parent generation, the expected mean values of the vector elements in 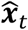 will be zero, i.e., 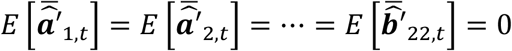. However, owing to different fitness (number of descendants) among the individuals in the parent generation, the corresponding mean values in the offspring generation before new reproduction will be different from 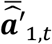 etc., and these within-generation differences may be used for updating of the mean reaction norm parameters 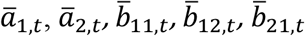 and 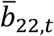 in Eq. (6a,b). After this updating the offspring are ready to become new parents, again with 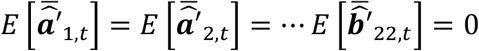.

Selection will thus result in within-generation incremental changes in the mean values of the predicted random effects from parents to offspring before reproduction, generally given by Eq. (1),

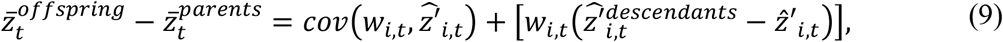

where 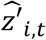 is any predicted individual value 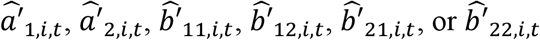 in the random effects vector 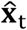 in Eq. (8), while 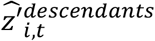 is the corresponding mean value for the descendants of individual *i* in generation *t*. In Eq. (9), 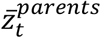 stands for the mean value of any one of the predicted random effects from Eq. (8), i.e., 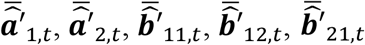, or 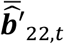, while 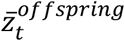 is the corresponding mean value for the offspring after selection but before reproduction. These incremental changes should thus at each generation be used for updating of the mean reaction norm parameter values before the offspring become new parents. For this purpose, we need an additional assumption, which follows from Assumption 3 above, because 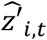 and 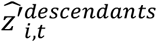 are predictions of an additive genetic component of a reaction norm parameter, and thus have no non-additive components:

#### Assumption 7

*There are no expected fitness weighted changes in predicted random effects from individual parents to their descendants after selection but before reproduction, i*.*e*., 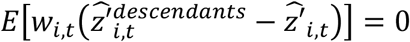.

When Assumption 7 is applied on Eq. (9), we obtain the Robertson-Price identity for the within-generation changes in the mean values (Ch. 6, Walsh and Lynch, 2018). Here, the incremental changes 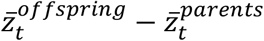 will be entirely determined by the additive genetic values 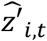 and individual fitness, and when these changes are used for updating we thus obtain between-generation changes in the mean values as given by Robertson’s secondary theorem of natural selection (Ch. 6, Walsh and Lynch, 2018). When Eq. (8) is applied on a given parent generation, and when the changes in mean values of predicted random effects from the parent to the offspring generation are used for updating, the incremental changes in those values under Assumption 7 thus follow from the following theorem:

#### Theorem 1

*In a population that is adequately described by Eq. (7), the incremental changes in mean reaction norm parameter values from generation to generation are found from Robertson*’*s secondary theorem of natural selection*,

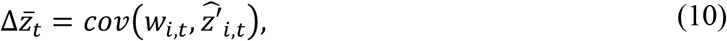

*where* 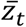 *is any mean parameter value* 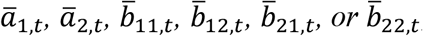, *while* 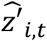 *is the corresponding predicted individual value* 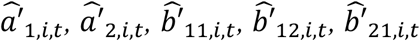, or 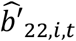, *in the random effects vector* 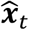 *in Eq. (8)*.

From Theorem 1 follows the incremental changes in mean traits according to

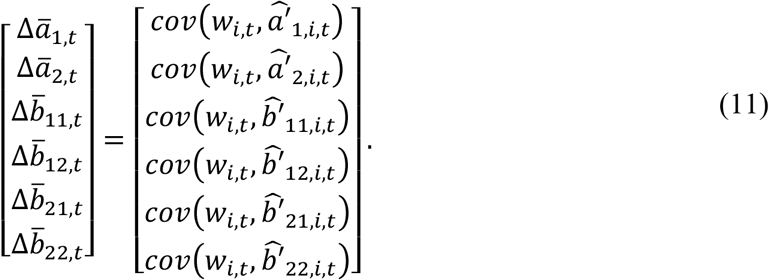

With correct initial mean reaction norm parameter values, this gives 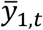 and 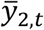 according to Eq. (6b), and this can be generalized to higher dimensions, with *p* > 2 and *q* > 2.

For finite population sizes there will be drift in the mean reaction norm parameter values, owing to random errors in the covariance computations according to Eq. (11). However, as we will see in the simulations, there are also other sources of drift.

From the BLUP theory above follows the following theorem:

#### Theorem 2

*For a reaction norm evolutionary system with p* = 2 *phenotypic traits and q* = 2 *environmental cues, Eq. (8) and Theorem 1 will with* ***A***_*t*_ = ***I***_*n*_ *result in incremental changes in mean reaction norm parameter values according to*

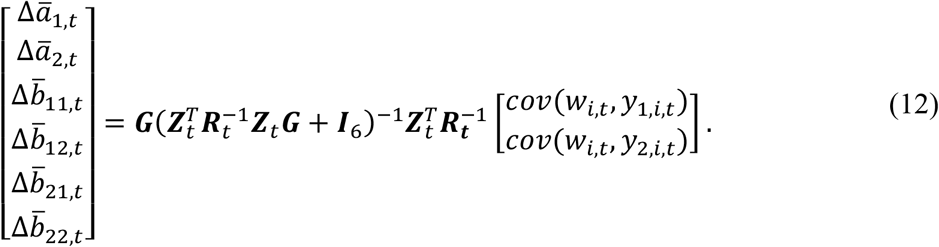

*This can be generalized to higher dimensions, with p* > 2 *and q* > 2.

See Appendix A for proof, and results in Section 3.

For comparisons with the multivariate breeder’s equation, the results from Eq. (12) can also be found by a generalization of the GRAD incremental parameter changes according to Eq. (5):

#### Theorem 3.

*Equation (12) can be reformulated as an extension of Eq. (5)*,

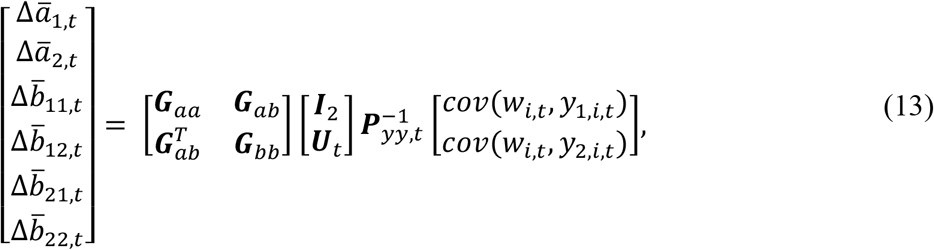

*where* 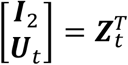, while 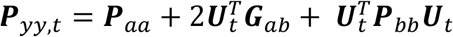.

See Appendix B for proof, and simulation results in Section 3, where Eqs. (12) and (13) give identical results for population sizes *n* ≥ 2. The simulations also show that the results from Eq. (13) are close to the results from a corresponding version of the multivariate breeder’s equation (4), with declining differences for increasing population size.

### 2.6 Example case without plasticity

When all plasticity slope parameter values are zero, Eq. (6a) gives 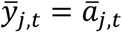 and the individual phenotypic traits

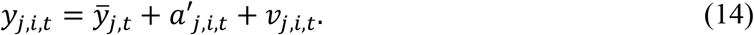

For *p* traits this leads to Eq. (8) with 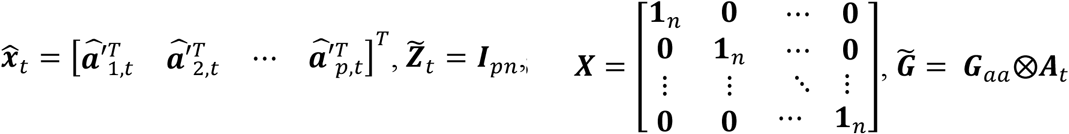 and 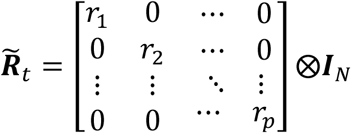, where *r* _*j*_ is the variance of υ_*j,i,t*_. For ***A***_*t*_ = ***I***_*n*_ this leads to the following theorem:

#### Theorem 4

*With individual phenotypic traits according to Eq. (14), and with* ***A***_*t*_ = ***I***_*n*_, *the dynamical BLUP model above and the multivariate breeder*’*s equation (2) give identical results*.

A proof of Theorem 4 is given in Ergon (2022c), although then relying on a comparison with the multivariate breeder’s equation. Here, Theorem 1 makes it into an independent proof.

Note that in this case Eq. (13) degenerates into Eq. (2).

### 2.7 Errors in estimated random effects variances

The fact that updated mean reaction norm parameter values, and thus also updated mean phenotypic trait values, are found by use of Robertson’s secondary theorem of natural selection applied on predicted random effects, as stated in Theorem 1, quite generally shows that the variances of these effects are underestimated. This is easily seen in the case without plasticity, given by Eq. (14), in which case Eq. (2) can be formulated as

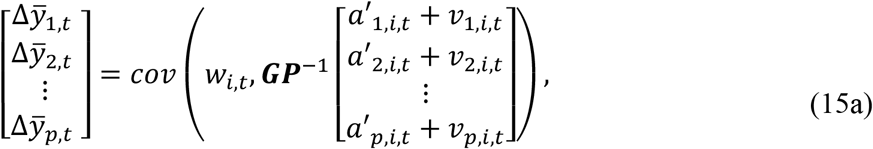

which should be compared with the result following from Theorem 1,

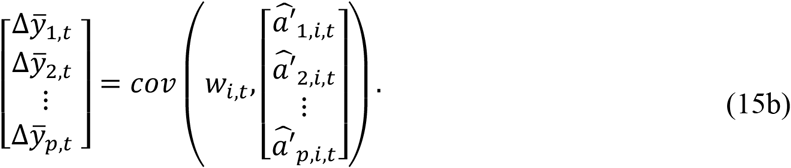

The underestimation of the variances 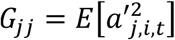 becomes especially transparent with diagonal ***G*** and ***P*** matrices, where we from Eqs. (15a,b), with use of 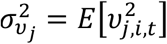, find 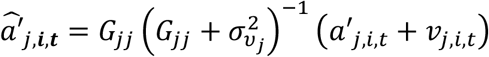. In this case we thus find

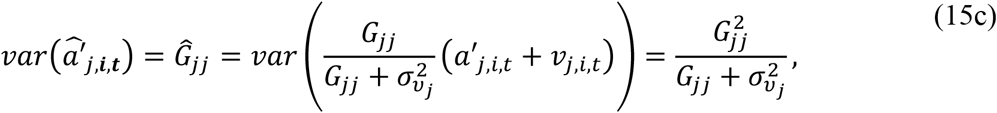

i.e., 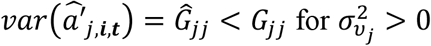. See Ergon (2022c) for simulation results.

2.8 Adjustments for overlapping generations

With surviving parents, only a fraction *f*_*t*_ < 1 of a given generation are offspring from the previous generation. The incremental changes in mean reaction norm parameter values from one generation to the next are then reduced accordingly, and by use of Eq. (11) we thus obtain

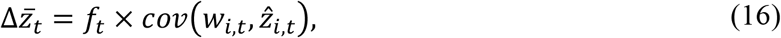

where 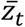 is any one of the mean reaction norm parameters, while 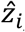 is the corresponding predicted individual random effect. As verified in the simulations, this will slow down responses on environmental changes, with reduced mean fitness as consequence.

## 3. Simulations

### 3.1 The aim of the simulations

The aim of the simulations is to verify the theoretical BLUP results by means of a toy example, and the purpose is fourfold. First, it is verified that mean reaction norm parameter values can be updated from generation to generation by means of Robertson’s secondary theorem of natural selection (Theorem 1). Second, it is shown that it is possible to disentangle the microevolutionary and plasticity components of for example climate change acclimations as shown in Eq. (11) in general, and in Eq. (12) for the special case with ***A***_*t*_ = ***I***_*n*_. Third, it is verified that the dynamical BLUP and GRAD results for the incremental changes in mean reaction norm parameter values are identical for population sizes *n* ≥ 2, provided that ***A***_*t*_ = ***I***_*n*_ (Theorems 2 and 3). Fourth, it is shown that the GRAD results are erroneous for populations with genetic relatedness between the individuals, i.e., for ***A***_*t*_ ≠ ***I***_*n*_.

### 3.2 Description of toy example

In a toy example in Ergon (2022a), the environmental input was a noisy positive trend in spring temperature, starting in 1970, and resulting in a noisy negative trend in mean breeding (clutch-initiation) date for a certain bird species, approximately as in fig. 2 in Bowers et al. (2016).

Here, the example is extended to include a second environmental variable with a noisy positive trend in mean value, and with variations from year to year that are somewhat positively correlated with the variations in spring temperature. This input may for example be a measure of spring rainfall. The example also includes a second adaptive phenotype, that might be the breeding habitat, as discussed in Chalfoun and Schmidt (2012). We thus have a microevolutionary system with two environmental cues and two phenotypic traits, similar to the theoretical example case.

The individual (mid-parent) fitness values are integers from 0 to 4, with number of descendants as unit, and cases with both non-overlapping and overlapping generations are simulated. The population size is assumed to be constant, which implies that not all descendants survive until reproduction. A constant population size is not essential for the principal results, but it simplifies the simulations.

In the simulations, the two environmental reference values are assumed to be known from historical data, i.e., it is assumed that the population was fully adapted to the stationary stochastic environment before the onset of anthropogenic and global climate change around 1970.

### 3.3 Environmental inputs and reaction norm model

Assume a population that is fully adapted to a stationary stochastic environment with mean spring temperature *u*_1,*ref*_ = 10 ℃ (the reference value for temperature), and mean spring rainfall *u*_2,*ref*_ = 2 mm/day (the reference value for rainfall). Also assume phenotypic scales such that the phenotypic values that maximize fitness in the reference environment are given by 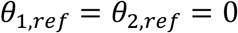. Further assume environmental cues 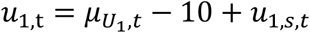, and 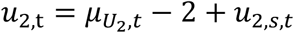, where the mean values 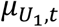 and 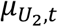 are ramp functions as shown in fig. 1, while *u*_1,*s,t*_ and *u*_2,*s,t*_ are zero mean and white random variables, i.e., without autocorrelation. In a corresponding way assume that 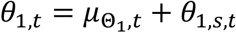 and 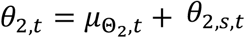, where 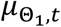 and 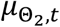 are ramp functions as shown in fig. 1, while *θ*_1,*s,t*_ and *θ*_2,*s,t*_ are zero mean and white random variables.

**Figure 1.**
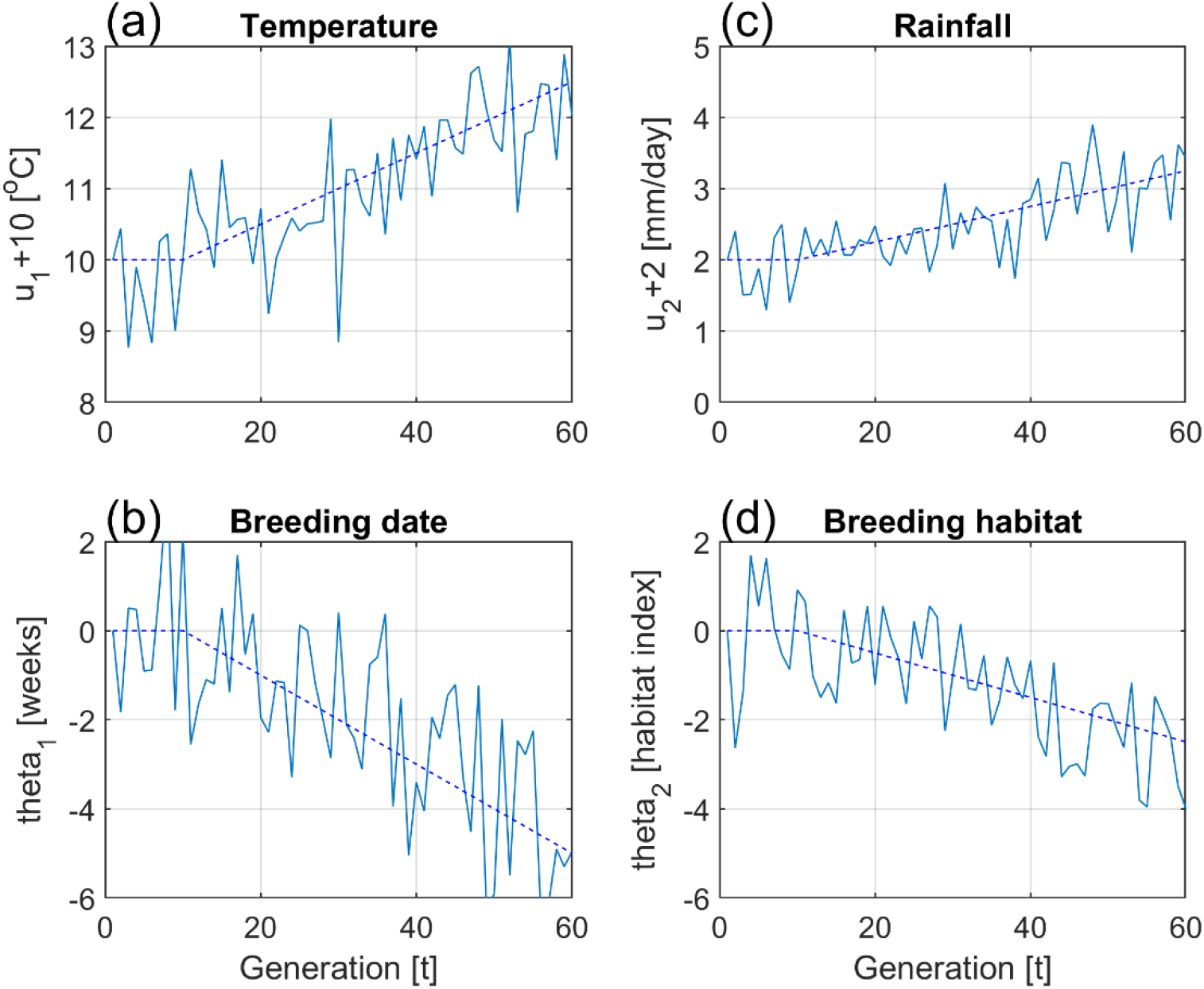
Typical input data for simulation example, with mean values shown by dashed lines, and with ramp functions starting at generation *t* = 10 (1970). Numerical values are 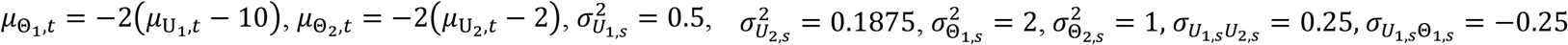, and 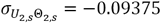.

Assume that *u*_1,*s,t*_, *u*_2,*s,t*_, *θ*_1,*s,t*_ and *θ*_2,*s,t*_ have a joint normal distribution with variances 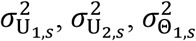 and 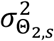, and covariances 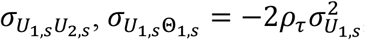 and 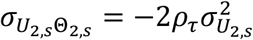, where *ρ*_*τ*_ is the autocorrelation of background environmental fluctuations, as described in more detail in Lande (2009). Data were generated for 60 generations, with typical input data as shown in fig. 1 (as mean values in breeding season). See Supplementary Material for MATLAB code.

Also assume an individual reaction norm model with two phenotypic traits, according to

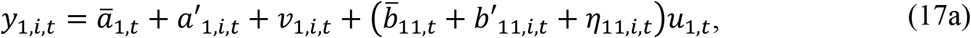

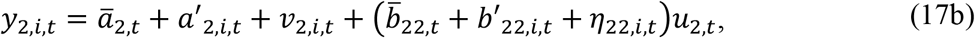

with parameters 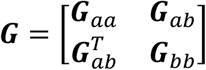 and 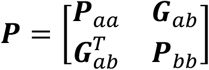, and parameter values 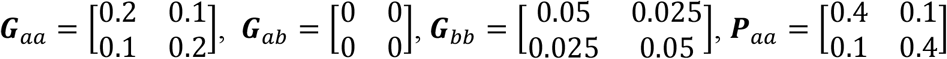 and 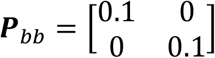. Note that Eqs. (17a,b) are somewhat simplified versions of Eq. (6a), in that 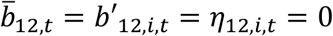, and 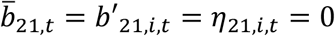. Also note that the two traits in Eqs. (17a,b) are correlated, with the covariance 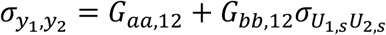.

### 3.4 Fitness function and initial mean reaction norm values

The individual fitness function is assumed to be rounded values of

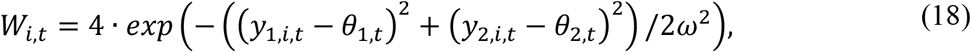

where *θ*_1,*t*_ and *θ*_2,*t*_ are the phenotypic values that maximize fitness, while *ω*^2^ = 10. The discrete values of *W*_*i,t*_ (number of descendants) are thus integers from 0 to 4.

In the simulations it is essential that the mean reaction norm parameters are given correct initial values at generation *t* = 1. We will assume that the phenotypic values are scaled such that the initial mean intercept values in a stationary stochastic environment are 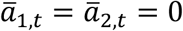, and that the initial mean reaction norm slope values are the optimal values in a stationary stochastic environment. These optimal values are the ones that maximize the expected individual fitness according to Eq. (18), in a stationary stochastic environment, and thus minimize the criterion functions 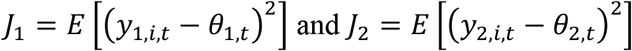 and 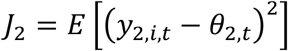. With 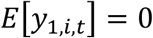, and substituting 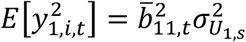 and 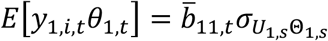, we find 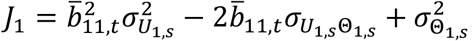, and setting 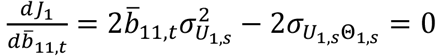, we thus find the optimal mean slope value 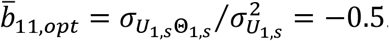. In the same way we find that 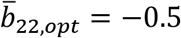 will minimize *J*_2_.

### 3.5 Simulation results

Simulation results with population size *n* = 100, and additive genetic relationship matrix ***A***_*t*_ = ***I***_*n*_, are shown in Fig. 2. New additive genetic (random) effects, and non-additive effects (residuals), were at each generation drawn from normal distributions in accordance with the given ***G*** and ***P*** matrices. The BLUP results given by Eq. (12) (green lines), and the GRAD results given by Eq. (13) (dashed blue lines), are identical for population sizes *n* ≥ 2. Results given by the multivariate breeder’s equation (4) (dotted magenta lines), are somewhat different from the BLUP results. These differences are clearly smaller for a population size of *n* = 1,000. Note that 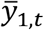 and 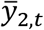 lag behind *θ*_1,*t*_ and *θ*_2,*t*_, as shown in Fig. 1, which is typical for ramp responses from dynamical systems with time constants.

**Figure 2.**
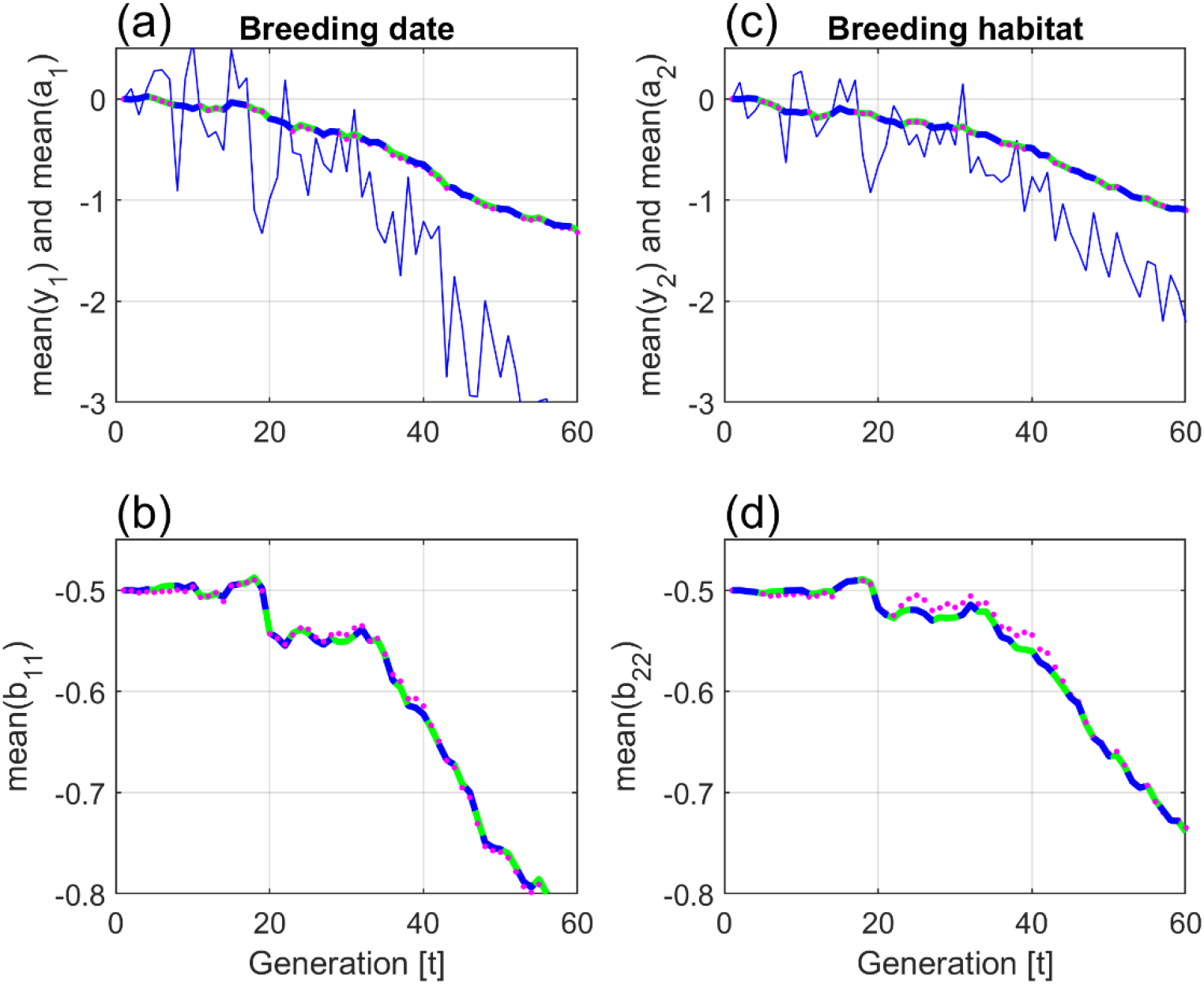
Simulation results with population size *n* = 100, and an additive genetic relationship matrix ***A***_*t*_ = ***I***_*n*_. Mean trait values 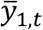 and 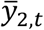 are shown by solid blue lines. Mean reaction norm parameter values are shown by solid green lines (BLUP), dashed blue lines (GRAD), and dotted magenta lines (the multivariate breeder’s equation).

As a test, the simulations were repeated with a constant additive genetic relationship matrix for a population with a high degree of relatedness among individuals, as shown for *n* = 6 in Eq. (19),

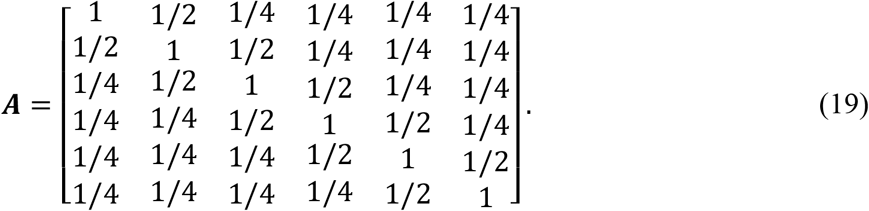

As shown in Fig. 3, this gave different results for the BLUP and GRAD methods. In this case new additive genetic (random) effects at each new generation were found as 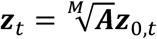, where ***z***_*t*_ stands for ***a***′_1,*t*_, ***a***′_2,*t*_, ***b***′_11,*t*_, ***b***′_12,*t*_, ***b***′_21,*t*_, or ***b***′_22,*t*_, and where the different data vectors ***z***_0,*t*_ were drawn from normal distributions in accordance with the given ***G*** matrix. Here, ^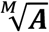^ is the matrix square root, i.e., 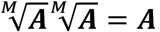. This reduced the variances of the random effects to approximately 75% of the nominal values. The random effects were also mean centred.

**Figure 3.**
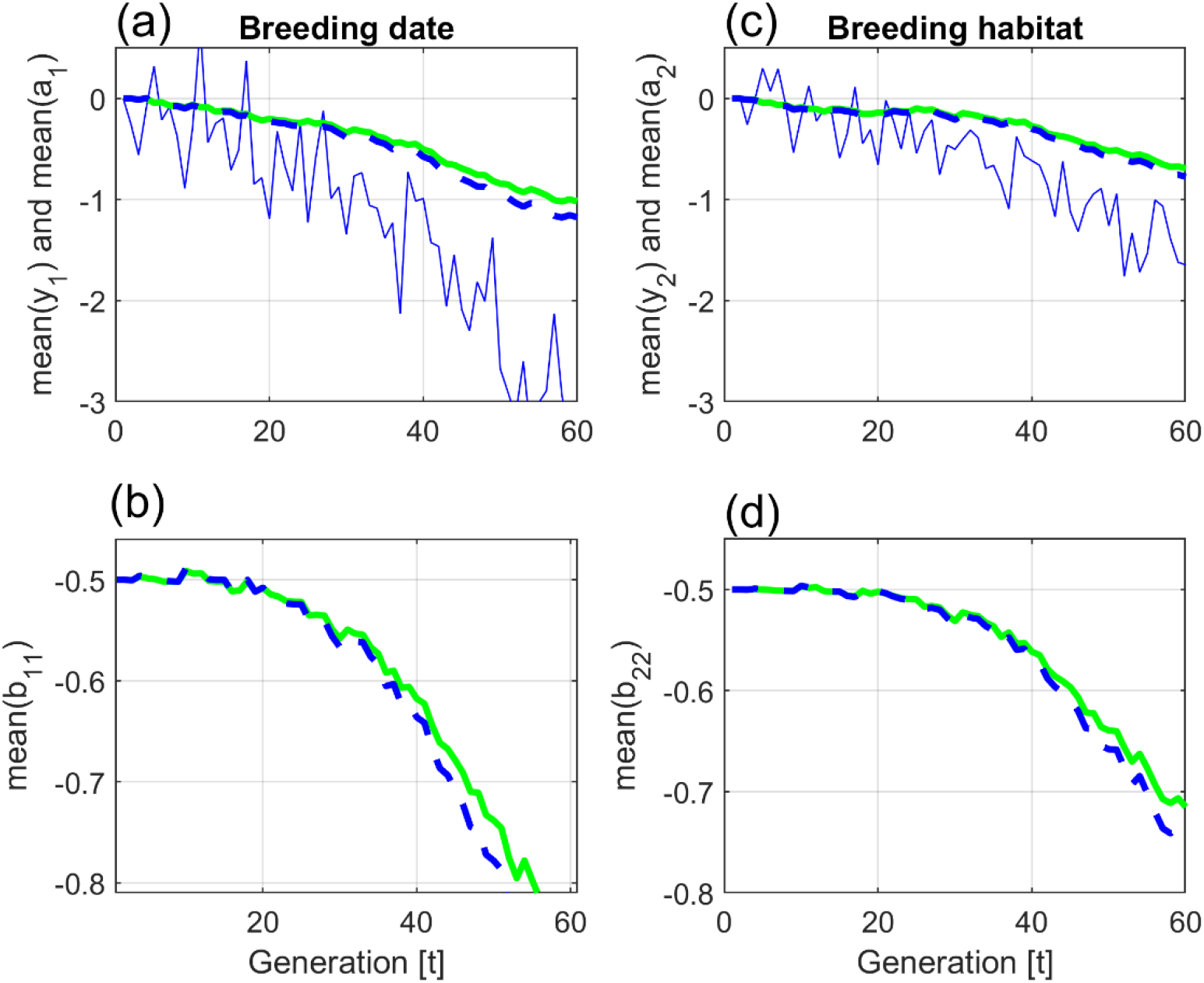
Simulation results with population size *n* = 100, and an additive genetic relationship matrix ***A*** according to Eq. (19). Mean trait values are shown by solid blue lines. Mean reaction norm parameter values are shown by solid green lines (BLUP) and dashed blue lines (GRAD).

The simulations with results as in Fig. 3 were repeated for a case with surviving parents, where only a fraction *f*_*t*_ = 0.5 of any given generation are offspring from the previous generation, i.e., with use of Eq. (16). As seen in Fig. 4, panels (a) and (c), and as must be expected, the responses are slowed down. As can also be seen in panels (a) and (c), the fraction *f*_*t*_ < 1 causes the mean trait values 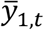 and 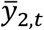 to lag further behind the phenotypic values *θ*_1,*t*_ and *θ*_2,*t*_ that maximize fitness. The result of this is lower mean fitness values, as can be seen in the (identical) plots in panels (b) and (d).

**Figure 4.**
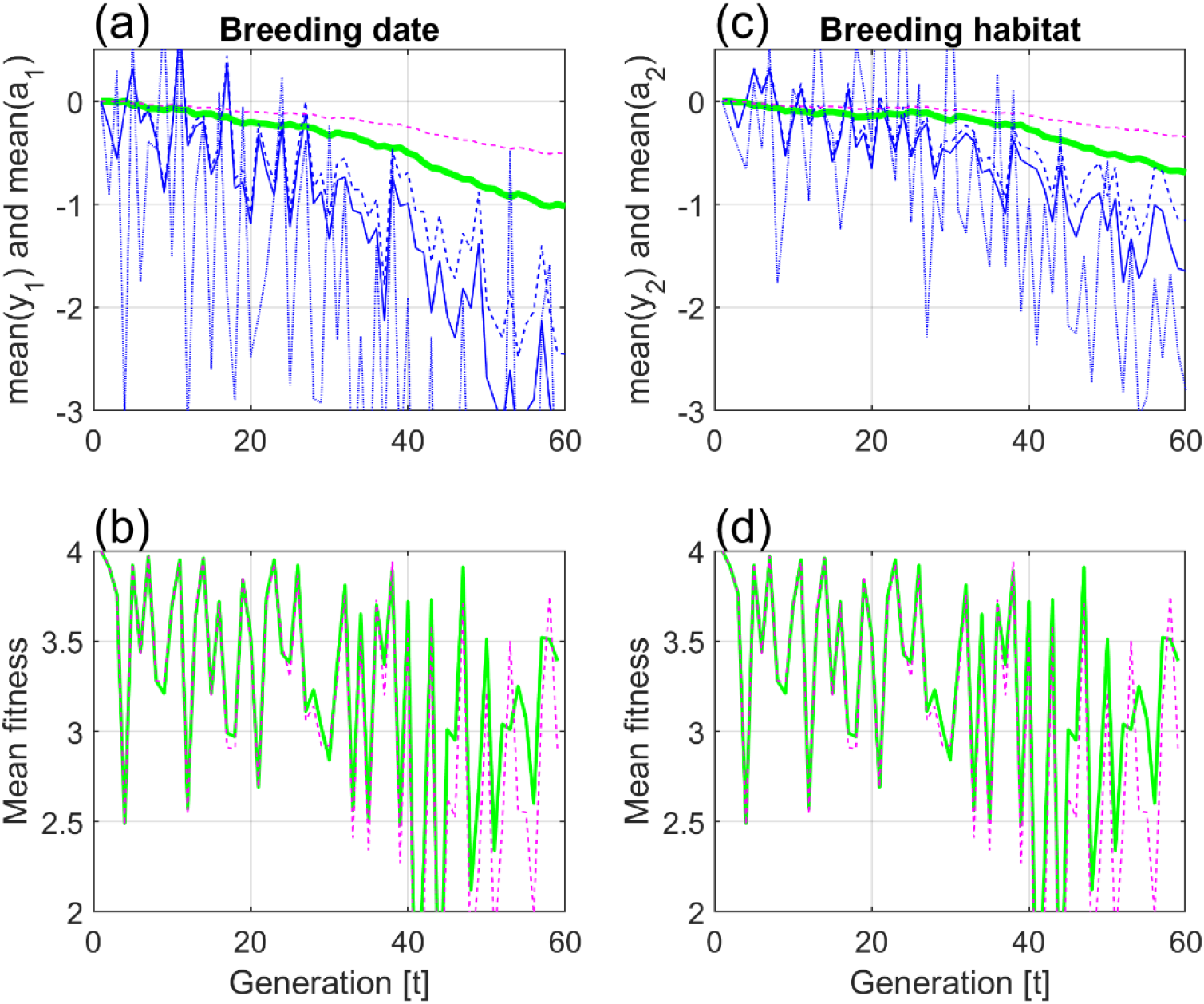
Simulation results with population size *n* = 100, and additive genetic relationship matrix ***A***_*t*_ ≠ ***I***_*n*_ according to Eq. (19). In panels (a) and (c), mean trait values 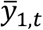 and 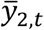 and mean reaction norm parameter values *ā*_1,*t*_ and *ā*_2,*t*_ for *f*_*t*_ = 1 are shown by solid blue and solid green lines, respectively. Dashed blue and magenta lines show the corresponding responses with a fraction *f*_*t*_ = 0.5 of new offspring in the population at all generations. Phenotypic values *θ*_1,*t*_ and *θ*_2,*t*_ that maximize fitness are added as weak dotted blue lines. Panels (b) and (d) show identical plots of mean fitness for *f*_*t*_ = 1 (solid green lines) and *f*_*t*_ = 0.5 (dashed magenta lines).

The changes in mean values over 60 generations in Figs. 2 and 3, will to some degree be caused by drift. As a test, the total change in *ā*_1,*t*_ over 60 generations in a stationary stochastic environment as before *t* = 10 in fig. 1, was computed. For population sizes from *n* = 10 to 400, and based on 100 repeated simulations this change had a mean value of approximately zero, and a standard error of approximately 0.1, i.e., around 10% of the corresponding changes in panel (a) in Figs. 2 and 3. With *n* = 2, this standard error due to drift increased to 15%. There were no noticeable differences between cases with ***A***_*t*_ ≠ ***I***_*n*_ and ***A***_*t*_ = ***I***_*n*_. In order to find an explanation of the drift, the environmental cue *u*_1,*t*_ over 60 generations in a stationary environment was approximated by a least squares straight line, denoted 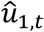 Based on 100 repeated simulations with *n* = 100, the resulting change in 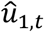 over 60 generations had a mean value of approximately zero, and a standard error of approximately 0.30, i.e., around 12% of the corresponding changes of *u*_1,*t*_ in Fig. 1, panel (a). This result indicates that the major part of the drift is caused by the stochastic nature of the environmental cues and the phenotypic values that maximize fitness.

## 4. Summary and Discussion

As shown in Section 2, a general reaction norm model with *p* phenotypic traits and *q* environmental cues, i.e., with *p*(1 + *q*) mean reaction norm parameters, can be formulated as a linear mixed model with fixed and random effects. In this model, the fixed effects will be the mean trait values, while the random effects will be the additive genetic components of individual deviations from mean reaction norm parameter values. From this follows that for any given parent generation, estimates of the mean trait values, and predictions of the additive genetic components of individual deviations from mean reaction norm parameter values, can be found from BLUP equations. Because the incidence and residual covariance matrices are functions of the environmental cues, and thus of time, I have introduced the concept of dynamical BLUP. Note that the elements in the random effects vector in Eq. (7) could be ordered differently, for example as 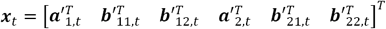 but then also the ***G, P, U***_*t*_ and **Z**_*t*_ matrices must be reorganized accordingly.

The development of the dynamical BLUP model in Eq. (8) relies on Assumptions 1, 2, 3, and 7, as given in Section 2, while Assumptions 4, 5, and 6 are needed only for the comparison with results based on the multivariate breeder’s equation. Most importantly, Assumption 4 implies that applications of the multivariate breeder’s equation on non-normal data in general will produce incorrect microevolutionary results. This problem was acknowledged by Lande and Arnold (1983), who in their Appendix proposed a possible correction for errors caused by skewness. The fundamental reason for such errors is that the multivariate breeder’s equation involves only mean and variance values of the additive genetic and non-additive effects, while influences from third and higher order statistical moments are ignored. See for example discussions in Bonamour et al. (2017) and Pick et al. (2022).

Although the development of the dynamical BLUP model in Eq. (8) does not rely on normal data (Robinson, 1991), it still follows from Theorems 2 and 3 that the BLUP results will be incorrect in cases with non-normal data and an additive genetic relationship matrix ***A***_*t*_ = ***I***_*n*_. A natural conclusion is therefore that non-normal data will give erroneous BLUP results also when ***A***_*t*_ ≠ ***I***_*n*_, at least for the dynamical BLUP model used here.

In the derivation of the linear mixed model in Eq. (7), I assumed linear reaction norms as functions of environmental cues *u*_1,*t*_, *u*_2,*t*_, etc., but there is nothing in the theory that prevents us from use of non-linear reaction norms that are also functions of 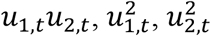, etc., as discussed in Gavrilets and Scheiner (1993) and Ergon (2018).

As shown by Theorem 1, updating of the mean reaction norm parameter values from generation to generation can be done by applying Robertson’s secondary theorem of natural selection on the vector elements 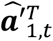, etc., in the predicted random effects. This shows that the well-known underestimation of the variances of the random effects, is just what is needed, in order to find the correct incremental changes in mean reaction norm parameter values. The resulting dynamics will depend on the additive genetic and residual covariance matrices ***G*** and ***R***_*t*_, as well as of the additive genetic relationship matrix ***A***_*t*_ for the parent generation.

The references to the Robertson-Price identity and Robertson’s secondary theorem of natural selection in connection with Assumption 7 and Theorem 1, might be somewhat confusing. In general (Ch. 6, Walsh and Lynch, 2018), the Robertson-Price identity follows from the Price equation, Eq. (1), by disregarding the second term on the righthand side. With *z*_*i,t*_ = *x*_*i,t*_ + *e*_*i,t*_ (Assumption 1), this leads to 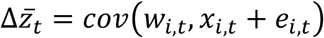. Robertson’s secondary theorem of natural selection, on the other hand, states that 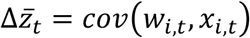, where *x*_*i,t*_ is the breeding value. For the special application of the Price equation in Eq. (9), we have 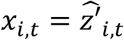 and *e*_*i,t*_ = 0, such that there is no difference between 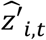 considered as a trait, and the breeding value of that trait. From this follows that Robertson-Price identity and Robertson’s secondary theorem of natural selection give the same results.

It is worth noticing that the additive genetic relationship matrix for the offspring generation does not affect the updating of mean reaction norm parameter values. This is shown in the theoretical derivations, but it is also a natural consequence of the fact that the dynamical BLUP model, just as the multivariate breeder’s equation, is an evolutionary state-space model (Ergon, 2018).

For cases with ***A***_*t*_ ≠ ***I***_*n*_, it should be noted that ***A***_*t*_ is included in the BLUP matrix equation (8) via 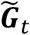 just as in the standard equation used in domestic breeding (Ch. 26, Lynch and Walsh, 1998). It is also reassuring to know that the BLUP equation asymptotically, for ***A***_*t*_ → ***I***_*n*_, gives results that are identical to the GRAD results based on the multivariate breeder’s equation (Theorems 2 and 3). Note that results from use of the multivariate breeder’s equation are equal to the GRAD results only asymptotically, i.e., for population size *n* → ∞. For cases without plasticity, and with ***A***_*t*_ = ***I***_*n*_, the results from all three methods are identical, independent of population size (Theorem 4).

Theorems 1, 2, and 3 were verified in simulations with use of ***A***_*t*_ = ***I***_*n*_, and population sizes down to *n* = 2, while Theorem 4 was verified by simulations in Ergon (2022c). Simulation results with ***A***_*t*_ ≠ ***I***_*n*_ were found by use of a constant and possibly unrealistic additive genetic relationship matrix, but these results still serve the purpose of showing that the BLUP and GRAD results with ***A***_*t*_ ≠ ***I***_*n*_ are different.

The changes in mean trait and mean reaction norm parameter values over 60 generations that can be seen in figs. 2 and 3, are partly caused by drift. As indicated at the end of Section 3, the major sources of drift are the stochastic nature of the input environmental variables *u*_1,*t*_ and *u*_2,*t*_, as well as of the phenotypic values *θ*_1,*t*_ and *θ*_2,*t*_ that maximize fitness. In the simulations, the population size plays a vital role only when *n* < 10.

In the simulations, the ***G*** and ***P*** parameters are assumed to be known, although they are normally not, and this is the case also for the reference environment and the initial mean reaction norm parameters. However, as shown in Ergon (2022a,b), these parameters can be found by a prediction error method (PEM) using all available input-output experimental or field data, including individual fitness data. Ergon (2022a,b) combined PEM with the GRAD model, i.e., with extensions of Eq. (5), but as will be reported separately, PEM works just as well when combined with the BLUP model in Eq. (8). BLUP/PEM will in fact be much better than GRAD/PEM when it comes to identification of the reference environment. As seen for GRAD/PEM in Ergon (2022a,b), and as will be reported separately for BLUP/PEM, the essential feature of PEM in the present setting is to predict the reaction norms as well as possible, while parameter values may be very much influenced by random measurement errors and modeling errors.

For the interested reader, the essential steps in the proposed dynamical BLUP method are summarized in Appendix C, where also the procedure for PEM system identification from laboratory or field data is included.

Parameter estimation by means of restricted maximum likelihood (REML) applied on data from a single generation is not possible for plastic organisms. A simple reason for this is that the variance components *r*_*j,t*_ in the diagonal residual covariance matrix ***R***_*t*_ are functions of several unknown variances of the non-additive effects in the reaction norm model, i.e., *r*_1,*t*_, *r*_2,*t*_, etc. can be found, but not all the residual covariances involved. The reason behind this problem is that there are confounding variables in the reaction norm equations. In for example Eq. (3), all the terms in *a*′_*i,t*_ + *b*′_*i,t*_*u*_*t*_ + *ν*_*i,t*_ + *η*_*i,t*_*u*_*t*_ are confounded, which implies that use of REML at a single generation will not give details of the ***G*** and ***R***_*t*_ matrices. As pointed out above, a solution is to use a prediction error method (PEM) where data from all generations are used. Note that this confounding is not a problem in the modeling process, where *ā*_*t*_ and 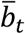 in 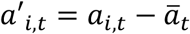 and 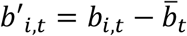 respectively, are found from Eq. (11), and where *a*′_*i,t*_, *b*′_*i,t*_, *ν*_*i,t*_ and *η*_*i,t*_ are samples with given variances.

With surviving parents, only a fraction *f*_*t*_ of the population will be offspring from the previous generation, and as shown in Eq. (16), the incremental changes in mean reaction norm parameter values will then be reduced accordingly. As verified in simulations, and as must be expected, the responses on environmental changes will then be slowed down, with reduced mean fitness as consequence. Note that the last term on the righthand side of Eq. (1), the Price equation, in general is affected by surviving parents, but that this term according to Assumption 7 does not affect the dynamical BLUP updating from generation to generation.

## Supporting information

MATLAB code

## Acknowledgements

The GRAD model in Eq. (5) was inspired by a comment by Michael B. Morrissay in a review of a quite different manuscript. I thank Bruce Walsh for constructive review comments, and Tormod Ådnøy for input regarding matrix details in BLUP modeling. I thank University of South-Eastern Norway for support and funding.

## Author contribution

Rolf Ergon is the sole author of this article.

## Data availability statement

MATLAB code for simulations is given in Supplementary Material archived on bioRxiv **doi: https://doi.org/10.1101/2023.04.09.536146doi:**.

## Appendix A Proof of Theorem 2

### Lemma 1

*Assuming an additive genetic relationship matrix* ***A***_*t*_ = ***I***_*n*_, *the mean values of the estimated fixed effects in Eq. (8), for j from* 1 *to p, are* 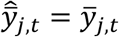 *while the mean values of the predicted random effects, for j from* 1 *to p and k from* 1 *to q, are* 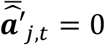 *and* 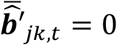.

***Proof:*** Eq. (8) will after elimination of 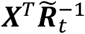 give the following equation for a specific trait *j*,

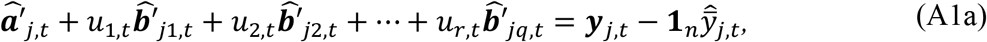

Taking mean values on both sides of Eq. (A1a) gives

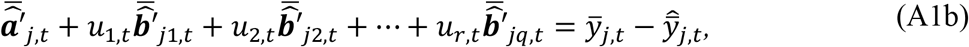

Eq. (8) also gives

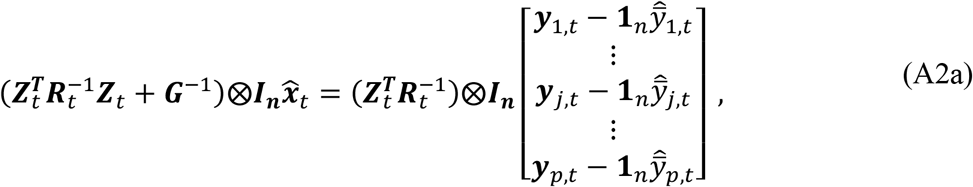

from which with use of the mixed model property (***A***⨂***B***)(***C***⨂***D***) = (***AC***)⨂(***BD***) follows that

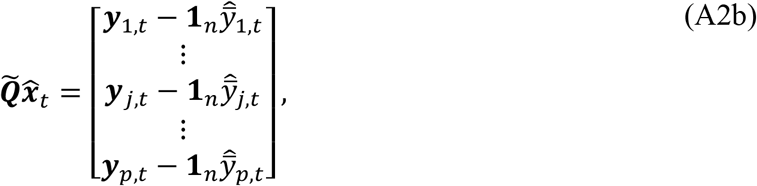

where 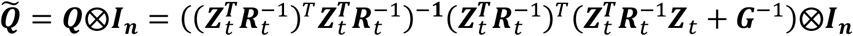 is an *pn* × *pn*(1 + *q*) matrix, with block row vectors 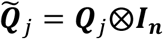 for *j* from 1 to *p*. From Eq. (A2b) thus follows

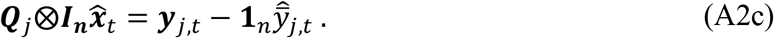

Since 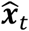 has dimension *n* × 1, we here have 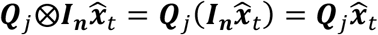 and by taking mean values of both sides of Eq. (A2c) we thus find

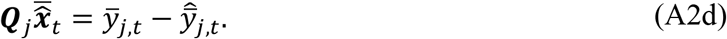

Eqs. (A1b) and (A2d) thus give two expressions for 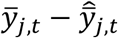, and since in general 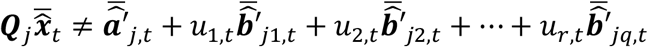, these expressions are equal if and only if 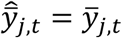 and 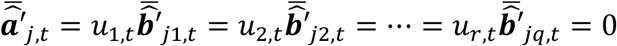.

Having established that ***A***_*t*_ = ***I***_*n*_ results in 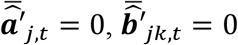 and 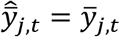 it is time to find how fitness affects the incremental changes in mean reaction norm parameter values. For clarity of presentation, we here limit the number of phenotypic traits to *p* = 2, and the number of environmental cues to *q* = 2, as we also did in parts of Section 2. We also introduce the diagonal *n* × *n* matrix ***F***_*t*_ = *diag*(*w*_*i,t*_), and the following two lemmas:

### Lemma 2

*The mean value of the product of* ***F***_*t*_ *and a predicted random effect vector* 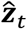 *in Eq. (8), where* 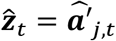 *for j from 1 to p, or* 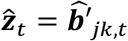 *for j from 1 to p and k from 1 to q, is* 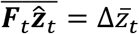 *i*.*e*., *the incremental change in the mean reaction norm parameter value* 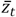.

***Proof:*** Since 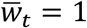 and 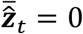 (from Lemma 1), we find 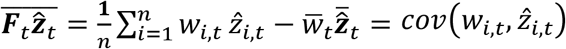 and thus according to Theorem 1, 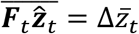.

### Lemma 3

*The mean value of the sum* 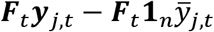 *for j from 1 to p, is cov*(*w*_*i,t*_, *y*_*j,i,t*_).

***Proof:*** 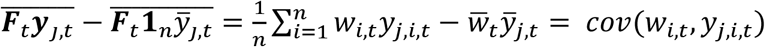.

Assuming *p* = 2 and *q* = 2, for simplicity, and using that 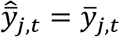 (Lemma 1), multiplication of Eq. (A2a) from the left with the block diagonal matrix 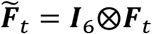, and insertion of 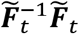 between 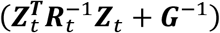 and 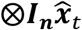 we find by use of the mixed model property (***A***⨂***B***)(***C***⨂***D***) = (***AC***)⨂(***BD***) that

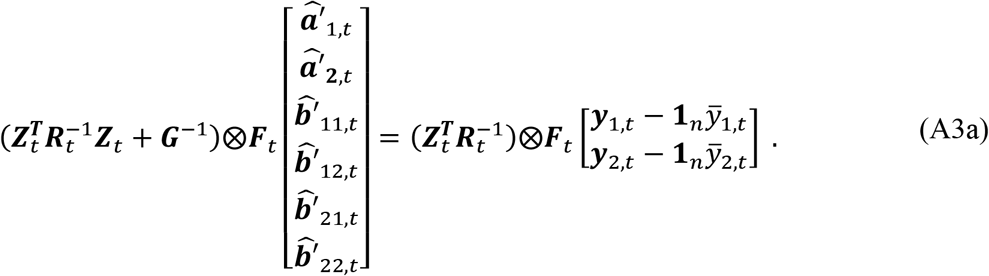

Since all elements in 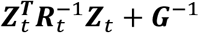 and 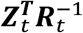 are multiplied by ***F***_*t*_, Eq. (A3a) can be reformulated as

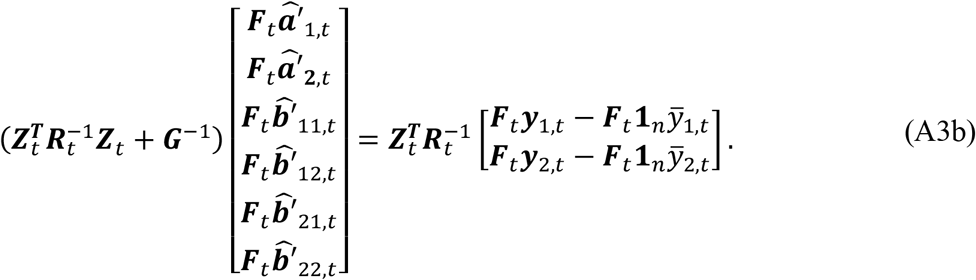

Taking mean values of all vector elements, will according to Lemma 2 and 3 finally give

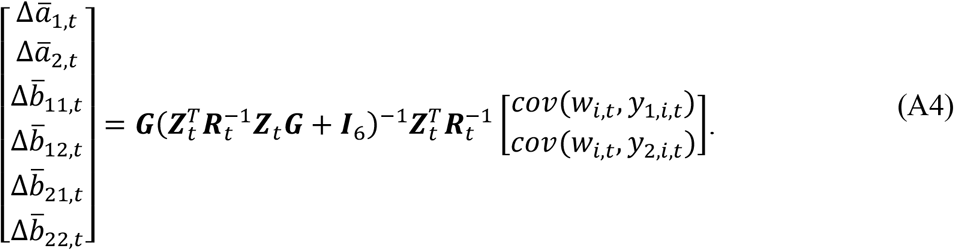

## Appendix B Proof of Theorem 3

Since 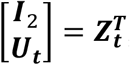, we must in order to prove Theorem 3 show that 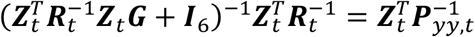. We can do that by use of the matrix inversion lemma (***A*** + ***BCD***)^−1^ = ***A***^−1^ − ***A***^−1^***B***(***C***^−1^ + ***DA***^−1^***B***)^−1^***DA***^−1^, with use of ***A*** = ***I***_6_, 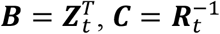 and ***D*** = **Z**_*t*_***G***. This gives

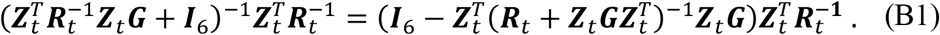

Here, 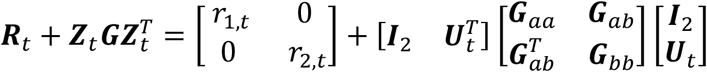, whitch with 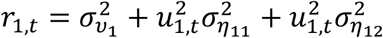 and 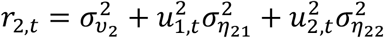 gives 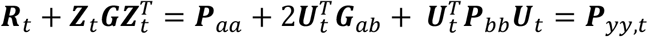, and thus 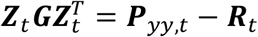. Eq. (B1) thus finally gives

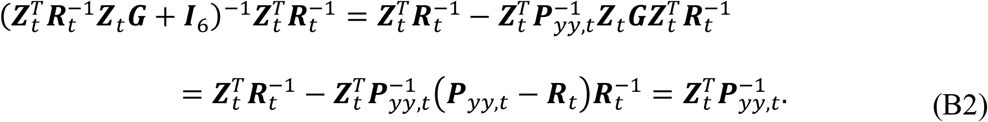

## Appendix C User guide, including PEM system identification

For the interested reader, I here summarize the essential steps in the proposed dynamical BLUP method, including the procedure for PEM system identification from laboratory or field data. Note that the feasibility of the PEM method in an evolutionary context has been tested in Ergon (2022a,b), but then with use of a GRAD model. For simplicity I here assume the BLUP simulation system in Section 3, with non-overlapping generations, and with input trends as shown in Fig. 1. With overlapping generations, the fraction *f*_*t*_ of the population that is offspring from the previous generation must be included as factors on the righthand sides of Eqs. (11) and (13).

1. Collect environmental input data *u*_1,*t*_ and *u*_2,*t*_, individual and mean phenotypic data 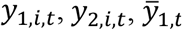 and 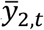, relative fitness data *w*_*i,t*_, and additive genetic relationship matrices ***A***_*t*_, for consecutive generations from *t* = 1 to *T*.
2. Form the dynamical incidence matrices 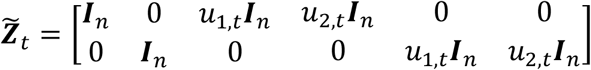.
3. 3.Set *G*_*aa*,11_ to an assumed and constant value (other ***G*** and ***P*** parameter values will be estimated relative to this value).
4. Assume some initial parameter values in 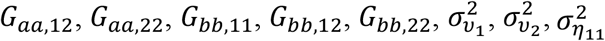 and 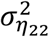. Simulations indicate that all these initial values may be set to zero. From this also follows initial parameter values in the dynamical residual covariance matrices 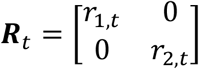, where 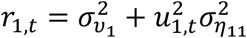 and 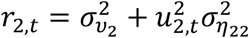.
5. Also assume initial values of the mean reaction norm slopes at time *t* = 1, i.e., 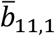 and 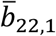, and of the reference environment values *u*_1,*ref*_ and *u*_2,*ref*_. Simulations indicate that 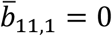 and 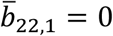 are useful values, while *u*_1,*ref*_ and *u*_2,*ref*_ preferably should be set to the mean values before the onset of the trends in Fig. 1.
6. Set initial values 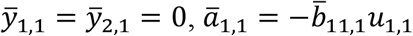, and 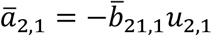, and predict the mean reaction norm parameter values for *t* = 1 to *T* by use of Eqs. (8) and (11). From this follow the predicted mean phenotypic values according to 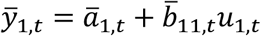 and 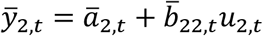.
7. Search for optimal parameter values by use of PEM, as shown in Fig (2) in Ergon (2022a), but with the dynamical BLUP model instead of a GRAD model. Use the function *fmincon* in MATLAB, or a corresponding function in, e.g., R.

A preliminary test with population size *n* = 100, and system identification by use of samples 41 to 60 in Fig. 1, gave good prediction results without measurement errors in the *u*_1,*t*_, *u*_2,*t*_, *y*_1,*i,t*_, *y*_2,*i,t*_ and *w*_*i,t*_ data. The optimization time for the BLUP optimization by use of an HP EliteBook × 360 1030 *G*3 laptop was as long as 3,000 seconds, owing to the repeated matrix inversions for computation of the predicted random effects in Eq. (8). No attempts were made to speed up the optimization by use of more efficient computations. Random measurement errors appear to affect especially the estimated reference environments, such that search bounds for these values should be rather narrow around the mean values of past stationary stochastic environments, which the population is judged to have been adapted to. Note that these reference values are not within the range of the input values from 41 to 60 in Fig. 1, which are used for the identification. Also note that for cases with ***A***_*t*_ = ***I***_*n*_, the GRAD predictions according to Eq. (13) give identical results, but with a very much shorter computation time.

