## Supplementary material for "Dynamical BLUP modeling of reaction norm evolution, accommodating changing environments, overlapping generations, and multivariate data": MATLAB code

for

Rolf Ergon

University of South-Eastern Norway

Porsgrunn, Norway

May 28, 2023

### MATLAB code

```
%-------------------------------------------------------------
% Dynamical BLUP modeling with Robertson-Price updatinng
% Rolf Ergon
% University of South-Eastern Norway
% March 4, 2023
%-------------------------------------------------------------

clear

%% Number of generations, population size, and width of fitness
function
T=60;
N=100;
wsquare=10;

%% G and P parameter values
Gaa11=0.2;
Gaa22=0.2;
Gaa12=0.1;
```

```
Gbb11=0.05;
Gbb22=0.05;
Gbb12=0.025;
varv11=Gaa11;
varv22=Gaa22;
vareta11=Gbb11;
vareta22=Gbb22;
Gaa=[Gaa11 Gaa12 ; Gaa12 Gaa22];
Gab=[0 0 ; 0 0 ];
Gbb=[Gbb11 Gbb12 ; Gbb12 Gbb22];
Paa=[Gaa11+varv11 Gaa12 ;  Gaa12 Gaa22+varv22];
Pbb=[Gbb11+vareta11 Gbb12 ; Gbb12 Gbb22+vareta22];
G=[Gaa Gab ; Gab' Gbb];
P=[Paa Gab ; Gab' Pbb];

%% Generate u and theta sequences for Fig. 1
vartheta1=2;
vartheta2=1;
varu1=0.5;
varu2=0.25;
rho=0.25;
u1plot=zeros(1,T);
u2plot=zeros(1,T);
du1=zeros(1,T);
dtheta1=zeros(1,T);
du2=zeros(1,T);
dtheta2=zeros(1,T);
for t=2:T
    du1(t)=sqrt(varu1)*randn;
    dtheta1(t)=du1(t)*rho*sqrt(vartheta1/varu1)+sqrt(vartheta1*(1-
rho^2))*randn;
    du2(t)=0.5*du1(t)+0.5*sqrt(varu2)*randn;
    dtheta2(t)=du2(t)*rho*sqrt(vartheta2/varu2)+sqrt(vartheta2*(1-
rho^2))*randn;
    if t>10
        u1plot(t)=(t-10)/20;
        u2plot(t)=(t-10)/40;
    end
end
u1=u1plot+du1;
u2=u2plot+du2;
theta1=-2*u1plot-dtheta1;
theta2=-2*u2plot-dtheta2;

%% Relationship matrix
for i=1:N
    for j=1:N
        if i==j dA(i,j)=0; end
        if i==j+1 dA(i,j)=0.5-0.25; end
        if i==j-1 dA(i,j)=0.5-0.25; end
    end
end
```

```
A=0.75*eye(N)+dA+0.25*ones(N,N);
A=eye(N);                          % Use for Fig. 2

%% Individual reaction norm parameter values around mean values
for t=1:T
    a1(:,t)=sqrt(Gaa11)*randn(N,1);
    a2(:,t)=Gaa12*a1(:,t)/Gaa11+sqrt(Gaa22-Gaa12^2/Gaa11)*randn(N,1);
    b11(:,t)=sqrt(Gbb11)*randn(N,1);
    b22(:,t)=Gbb12*b11(:,t)/Gbb11+sqrt(Gbb22-Gbb12^2/Gbb11)*randn(N,1);
    v1(:,t)=sqrt(varv11)*randn(N,1);
    v2(:,t)=sqrt(varv22)*randn(N,1);
    eta11(:,t)=sqrt(vareta11)*randn(N,1);
    eta22(:,t)=sqrt(vareta22)*randn(N,1);
    a1(:,t)=a1(:,t)-mean(a1(:,t));
    a2(:,t)=a2(:,t)-mean(a2(:,t));
    a1(:,t)=sqrtm(A)*a1(:,t);
    a2(:,t)=sqrtm(A)*a2(:,t);
    b11(:,t)=sqrtm(A)*b11(:,t);
    b22(:,t)=sqrtm(A)*b22(:,t);
    a1(:,t)=a1(:,t)-mean(a1(:,t));
    a2(:,t)=a2(:,t)-mean(a2(:,t));
    b11(:,t)=b11(:,t)-mean(b11(:,t));
    b22(:,t)=b22(:,t)-mean(b22(:,t));
    v1(:,t)=v1(:,t)-mean(v1(:,t));
    v2(:,t)=v2(:,t)-mean(v2(:,t));
    eta11(:,t)=eta11(:,t)-mean(eta11(:,t));
    eta22(:,t)=eta22(:,t) -mean(eta22(:,t));
end

%% BLUP predictions
abar1B=0*ones(1,T);
abar2B=0*ones(1,T);
bbar11B=-0.5*ones(1,T);
bbar22B=-0.5*ones(1,T);
ybar1B=0*ones(1,T);
ybar2B=0*ones(1,T);
for t=1:T-1
    bterm1=(bbar11B(t)+b11(:,t)+eta11(:,t)).*u1(t);
    bterm2=(bbar22B(t)+b22(:,t)+eta22(:,t)).*u2(t);
    y1(:,t)=abar1B(t)+a1(:,t)+v1(:,t)+bterm1;
    y2(:,t)=abar2B(t)+a2(:,t)+v2(:,t)+bterm2;
    W(:,t)=round(4*exp(-((y1(:,t)-theta1(t)).^2+(y2(:,t)-
theta2(t)).^2)/(2*wsquare)));
    Wbar(t)=mean(W(:,t));
    X=[ones(N,1) zeros(N,1); zeros(N,1) ones(N,1)];
    r1=varv11+vareta11.*u1(t)^2;
    r2=varv22+vareta22.*u2(t)^2;
    R=[r1 0 ; 0 r2];
    U=[u1(t) 0 ; 0  u2(t)];
    Z=[eye(2) U'];
    Ztilde=kron(Z,eye(N));
    Gtilde=kron(G,A);
```

```
    Rtilde=kron(R,eye(N));
    M=[X'*inv(Rtilde)*X X'*inv(Rtilde)*Ztilde
        Ztilde'*inv(Rtilde)*X Ztilde'*inv(Rtilde)*Ztilde+inv(Gtilde)];
    y=[y1(:,t) ; y2(:,t)];
    Ym=[X'*inv(Rtilde)*y ; Ztilde'*inv(Rtilde)*y];
    effects=inv(M)*Ym;
    cova1=cov(W(:,t),effects(3:N+2));
    Dabar1=((N-1)/N)*cova1(1,2)/Wbar(t);
    cova2=cov(W(:,t),effects(N+3:2*N+2));
    Dabar2=((N-1)/N)*cova2(1,2)/Wbar(t);
    covb11=cov(W(:,t),effects(2*N+3:3*N+2));
    Dbbar11=((N-1)/N)*covb11(1,2)/Wbar(t);
    covb22=cov(W(:,t),effects(3*N+3:4*N+2));
    Dbbar22=((N-1)/N)*covb22(1,2)/Wbar(t);
    abar1B(t+1)=abar1B(t)+Dabar1;
    abar2B(t+1)=abar2B(t)+Dabar2;
    bbar11B(t+1)=bbar11B(t)+Dbbar11;
    bbar22B(t+1)=bbar22B(t)+Dbbar22;
    ybar1B(t+1)=abar1B(t+1)+bbar11B(t+1)*u1(t+1);
    ybar2B(t+1)=abar2B(t+1)+bbar22B(t+1)*u2(t+1);
end

%% GRAD predictions
abarG=zeros(2,T);
bbarG=[ -0.5*ones(1,T)
        -0.5*ones(1,T) ];
bbar11G(1,1)=-0.5;
bbar22G(1,1)=-0.5;
for t=1:T-1
    U=[u1(t) 0 ;  0 u2(t)];
    Pyy=Paa+U'*Pbb*U;
    covWy1=(N-1)*cov(W(:,t),y1(:,t))/N;
    covWy2=(N-1)*cov(W(:,t),y2(:,t))/N;
    betay(:,t)=inv(Pyy)*[covWy1(1,2) ; covWy2(1,2)]/Wbar(t);
    abarG(:,t+1)=abarG(:,t)+Gaa*betay(:,t);
    bbarG(:,t+1)=bbarG(:,t)+Gbb*U*betay(:,t);
    abar1G(1,t+1)=abarG(1,t+1);
    abar2G(1,t+1)=abarG(2,t+1);
    bbar11G(1,t+1)=bbarG(1,t+1);
    bbar22G(1,t+1)=bbarG(2,t+1);
end

%% Predictions from the multivariate breeder's equation
abar1L=zeros(1,T);
abar2L=zeros(1,T);
bbar11L=-0.5*ones(1,T);
bbar22L=-0.5*ones(1,T);
ybar1L=zeros(1,T);
ybar2L=zeros(1,T);
xbar=zeros(4,T);
for t=1:T-1
    covW1=(N-1)*cov([a1(:,t)+v1(:,t) b11(:,t)+eta11(:,t) W(:,t)])/N;
```

```
    covW2=(N-1)*cov([a2(:,t)+v2(:,t) b22(:,t)+eta22(:,t) W(:,t)])/N;
    xbar(:,t)=[abar1L(t) abar2L(t) bbar11L(t) bbar22L(t)]';
    covxW=[covW1(1,3) covW2(1,3) covW1(2,3) covW2(2,3)];
    xbar(:,t+1)=xbar(:,t)+G*pinv(P)*covxW'/Wbar(t);
    xbarny=xbar(:,t+1);
    abar1L(t+1)=xbarny(1);
    bbar11L(t+1)=xbarny(3);
    abar2L(t+1)=xbarny(2);
    bbar22L(t+1)=xbarny(4);
    ybar1L(t+1)=abar1L(t+1)+bbar11L(t+1)*u1(t+1);
    ybar2L(t+1)=abar2L(t+1)+bbar22L(t+1)*u2(t+1);
end

%% Figure 1
figure(1)

subplot(2,2,1)
plot(u1+10), hold on
plot(u1plot+10,'--b'), hold off
axis([0 60 8 13]), grid
title('Temperature')
ylabel('u_1+10 [^oC]')
text(0,13.4,'(a)','FontSize',14)

subplot(2,2,2)
plot(u2+2), hold on
plot(u2plot+2,'--b'), hold off
axis([0 60 0 5]), grid
title('Rainfall')
ylabel('u_2+2 [mm/day]')
text(0,5.4,'(c)','FontSize',14)

subplot(2,2,3)
plot(theta1), hold on
plot(-2*u1plot,'--b'), hold off
axis([0 60 -6 2]), grid
title('Breeding date')
xlabel('Generation [t]')
ylabel('theta_1 [weeks]')
text(0,2.6,'(b)','FontSize',14)

subplot(2,2,4)
plot(theta2), hold on
plot(-2*u2plot,'--b'), hold off
axis([0 60 -6 2]), grid
title('Breeding habitat')
xlabel('Generation [t]')
ylabel('theta_2 [habitat index]')
text(0,2.6,'(d)','FontSize',14)

%% Figures 2 and 3
figure(2)
```

```
subplot(2,2,1)
plot(ybar1B,'b'), hold on
plot(abar1B,'g','LineWidth',2)
plot(abar1L,'.m','LineWidth',2)                    % Don't use for Fig. 3
plot(abar1G,'--b','LineWidth',2)
hold off, grid
title('Breeding date')
ylabel('mean(y_1) and mean(a_1)')
axis([0 T -3 0.5])
text(0,0.8,'(a)','FontSize',14)

subplot(2,2,3)
plot(bbar11B,'g','LineWidth',2), hold on
plot(bbar11G,'--b','LineWidth',2)
plot(bbar11L,'.m','LineWidth',2)                    % Don't use for Fig. 3
hold off, grid
xlabel('Generation [t]')
ylabel('mean(b_1_1)')
axis([0 T -0.8 -0.45])
text(0,-0.42,'(b)','FontSize',14)

subplot(2,2,2)
plot(ybar2B,'b'), hold on
plot(abar2B,'g','LineWidth',2)
plot(abar2L,'.m','LineWidth',2)                    % Dont use for Fig. 3
plot(abar2G,'--b','LineWidth',2)
hold off, grid
title('Breeding habitat')
ylabel('mean(y_2) and mean(a_2)')
axis([0 T -3 0.5])
text(0,0.8,'(c)','FontSize',14)

subplot(2,2,4)
plot(bbar22B,'g','LineWidth',2), hold on
plot(bbar22L,'.m','LineWidth',2)                   % Dont use for Fig. 3
plot(bbar22G,'--b','LineWidth',2)
hold off, grid
xlabel('Generation [t]')
ylabel('mean(b_2_2)')
axis([0 T -0.8 -0.45])
text(0,-0.42,'(d)','FontSize',14)


%% BLUP predictions wirh surviving parents
f=0.5;
abar1=0*ones(1,T);
abar2=0*ones(1,T);
bbar11=-0.5*ones(1,T);
bbar22=-0.5*ones(1,T);
ybar1=0*ones(1,T);
ybar2=0*ones(1,T);
```

```
for t=1:T-1
    bterm1=(bbar11(t)+b11(:,t)+eta11(:,t)).*u1(t);
    bterm2=(bbar22(t)+b22(:,t)+eta22(:,t)).*u2(t);
    y1(:,t)=abar1(t)+a1(:,t)+v1(:,t)+bterm1;
    y2(:,t)=abar2(t)+a2(:,t)+v2(:,t)+bterm2;
    Wc(:,t)=round(4*exp(-((y1(:,t)-theta1(t)).^2+(y2(:,t)-
theta2(t)).^2)/(2*wsquare)));
    Wbarc(t)=mean(Wc(:,t));
    X=[ones(N,1) zeros(N,1); zeros(N,1) ones(N,1)];
    r1=varv11+vareta11.*u1(t)^2;
    r2=varv22+vareta22.*u2(t)^2;
    R=[r1 0 ; 0 r2];
    U=[u1(t) 0 ; 0  u2(t)];
    Z=[eye(2) U'];
    Ztilde=kron(Z,eye(N));
    Gtilde=kron(G,A);
    Rtilde=kron(R,eye(N));
    M=[X'*inv(Rtilde)*X X'*inv(Rtilde)*Ztilde
        Ztilde'*inv(Rtilde)*X Ztilde'*inv(Rtilde)*Ztilde+inv(Gtilde)];
    y=[y1(:,t) ; y2(:,t)];
    Ym=[X'*inv(Rtilde)*y ; Ztilde'*inv(Rtilde)*y];
    effects=inv(M)*Ym;
    ybar1Btest(t)=effects(1);
    ybar2Btest(t)=effects(2);
    cova1=cov(W(:,t),effects(3:N+2));
    Dabar1=f*((N-1)/N)*cova1(1,2)/Wbar(t);
    cova2=cov(W(:,t),effects(N+3:2*N+2));
    Dabar2=f*((N-1)/N)*cova2(1,2)/Wbar(t);
    covb11=cov(W(:,t),effects(2*N+3:3*N+2));
    Dbbar11=f*((N-1)/N)*covb11(1,2)/Wbar(t);
    covb22=cov(W(:,t),effects(3*N+3:4*N+2));
    Dbbar22=f*((N-1)/N)*covb22(1,2)/Wbar(t);
    abar1(t+1)=abar1(t)+Dabar1;
    abar2(t+1)=abar2(t)+Dabar2;
    bbar11(t+1)=bbar11(t)+Dbbar11;
    bbar22(t+1)=bbar22(t)+Dbbar22;
    ybar1(t+1)=abar1(t+1)+bbar11(t+1)*u1(t+1);
    ybar2(t+1)=abar2(t+1)+bbar22(t+1)*u2(t+1);
end

%% Fig. 4
figure(4)

subplot(2,2,1)
plot(ybar1B,'b'), hold on
plot(abar1B,'g','LineWidth',2)
plot(ybar1,'--b')
plot(abar1,'--m')
plot(theta1,':b'), hold off, grid
title('Breeding date')
ylabel('mean(y_1) and mean(a_1)')
axis([0 T -3 0.5])
```

```
text(0,0.8,'(a)','FontSize',14)

subplot(2,2,2)
plot(ybar2B,'b'), hold on
plot(abar2B,'g','LineWidth',2)
plot(ybar2,'--b')
plot(abar2,'--m')
plot(theta2,':b'), hold off, grid
title('Breeding habitat')
ylabel('mean(y_2) and mean(a_2)')
axis([0 T -3 0.5])
text(0,0.8,'(c)','FontSize',14)

subplot(2,2,3)
plot(Wbar,'g','LineWidth',1), hold on
plot(Wbarc,'--m'), hold off, grid
xlabel('Generation [t]')
ylabel('Mean fitness')
axis([0 T 2 4])
text(0,4.15,'(b)','FontSize',14)

subplot(2,2,4)
plot(Wbar,'g','LineWidth',1), hold on
plot(Wbarc,'--m'), hold off, grid
xlabel('Generation [t]')
ylabel('Mean fitness')
axis([0 T 2 4])
text(0,4.15,'(d)','FontSize',14)
```
